# CGRP-receptor family reveals endogenous GPCR agonist bias and its significance in primary human cardiovascular cells

**DOI:** 10.1101/2020.12.21.423730

**Authors:** Ashley J. Clark, Niamh Mullooly, Dewi Safitri, Matthew Harris, Tessa de Vries, Antoinette Maassen Van Den Brink, David R. Poyner, Davide Giani, Mark Wigglesworth, Graham Ladds

**Affiliations:** Department of Pharmacology, University of Cambridge, Tennis Court Road, Cambridge, CB2 1PD, UK; Functional Genomics, Discovery Sciences, AstraZeneca, Cambridge CB2 0SL, United Kingdom; School of Life and Health Sciences, Aston University, Aston Triangle, Birmingham B4 7ET, United Kingdom; Department of Internal Medicine, Erasmus MC, Erasmus University Medical Centre, Rotterdam, Rotterdam, Netherlands; Hit Discovery, Discovery Sciences, BioPharmaceuticals R&D, AstraZeneca, Alderley Park SK10 4TG, UK

**Author notes:** **Address for correspondence**: Dr Graham Ladds, Department of Pharmacology, University of Cambridge, Tennis Court Road, Cambridge, CB2 1PD. **Author Contributions**: DRP, AMD, MW and GL conceived and designed the research; AJC, TV and DS performed the experiments; AJC, NM and DG designed the CRISPR-Cas9 experiments, AJC, GL, MH and MW analysed data; AJC, MH and GL wrote manuscript, DRP, AMVDB, MW and DG revised and edited the manuscript.

**Keywords:** G protein-coupled receptor, CGRP, adrenomedullin, adrenomedullin 2, receptor activity-modifying protein, agonist bias, gene editing, physiological bias

## Abstract

Agonist bias at G protein-coupled receptors has attracted considerable interest, although its relevance for physiologically-produced agonists is not always clear. Here, using primary human cells and gene editing techniques, we demonstrate for the first time, endogenous agonist bias with physiological consequences for the calcitonin-like receptor (CLR). We reveal that by switching the accessory protein: receptor activity-modifying protein (RAMP) associated with CLR we can re-route the physiological pathways activated by the stimulating peptide agonists. These results have revealed a unique role in calcium-mediated nitric oxide signalling for the little-understood peptide adrenomedullin 2 and distinct pro-proliferative effects of calcitonin-gene related peptide (CGRP) and adrenomedullin in cardiovascular cells. This work reveals that CLR-based agonist bias occurs naturally in human cells and has a fundamental purpose for its existence. We anticipate this will be a starting point for more studies into RAMP function in native environments and its importance in endogenous GPCR signalling.

## Introduction

G protein-coupled receptors (GPCRs) form the largest protein family in the human genome. ∼30% of marketed drugs target these receptors and therefore understanding their signalling pathways is not simply an academic exercise. For many years it had been incorrectly assumed that agonist-occupied GPCRs signalled through a single pathway to elicit their response. However, there is now overwhelming evidence to suggest that many GPCRs exist in multiple receptor conformations and can elicit numerous functional responses, both G protein- and non-G protein-dependent. Furthermore, different agonists, acting at the same receptor have the potential to activate different signalling pathways to varying extents; a concept referred to as biased agonism or signalling bias^1,2^. While the therapeutic promise of biased agonists is obvious^3^: it allows design of ligands that actively engage with one beneficial signalling outcome while reducing the contribution from those that mediate more undesirable effects, it is not without controversy. For example, recent doubt has been cast on validity of developing synthetic biased agonists against the μ-opioid receptor – a GPCR that has been considered the trailblazer for therapeutic potential of biased agonism^4^. Thus, further investigations into the role of agonist bias and its physiological importance, particularly its relevance to endogenous agonists, are required to bridge the gap between heterologous studies and in-vivo investigations.

While there are many well-studied GPCRs that exhibit signalling bias, including adrenoceptors, and the aforementioned μ-opioid receptor, we have chosen to focus upon the calcitonin-like receptor (CLR). Like many other GPCRs, CLR can couple to multiple G proteins and β-arrestins. Importantly, when co-expressed with one of three receptor activity modifying proteins, (RAMPs, see below), it can be activated by distinct endogenous agonists; calcitonin-gene related polypeptide (CGRP), adrenomedullin (AM) and adrenomedullin 2/intermedin (AM2). This makes it a good system to investigate the role of bias for such endogenous ligands. CGRP, an abundant neuropeptide, is the most potent microvascular vasodilator known. While it is thought to be cardioprotective, it has also been implicated in diseases such as migraine^5^. AM is released by the vascular endothelium and is also a potent vasodilator that can modulate vascular tone, it is involved in angiogenesis, and is elevated in some cancers and heart failure^6-8^. AM2 is also a vasodilator and highly expressed in the heart and vasculature^9,10^. It can cause sympathetic activation, have antidiuretic effects, and is upregulated in cardiac hypertrophy and myocardial infarction^11^.

Molecularly, CLR and its close relative, the calcitonin receptor (CTR), are classical class B GPCRs. CLR is pleiotropically coupled, predominately activating Gαs although there are reports of couplings to both G_i/o_ and G_q/11/142,12_ families. These Gα subunits promote activation/inhibition of adenylyl cyclase and phospholipase C to generate intracellular second messengers including cAMP and mobilise intracellular Ca^2+^ (Ca^2+^_i_) which then activate their respective intracellular signalling cascades. Beyond the Gα subunits CLR has been reported to couple to β-arrestins^13-13^ inducing internalisation, although it has been suggested that this interaction can also lead to its own signalling events possible promoting cell growth and proliferation^14^. Despite this high potential for agonist-induced pleiotropy, CLR remains most closely associated with adenylyl cyclase activation and generation of cAMP. It is unknown whether this is representative of CLR’s true signalling pattern in the endogenous setting.

Additional complexity is added to the pharmacology of the CLR since it has an absolute requirement for the formation of a heterodimer with a RAMP^15,16^. In overexpression studies, each of the three RAMPs have been shown to differentially influence the affinity and agonist bias of the CGRP family of peptides at the CLR^17,18^. CLR in complex with RAMP1 generates the CGRP receptor since CGRP has been demonstrated to be the most potent of the three agonists at this receptor for generation of cAMP. Likewise, CLR-RAMP2 generates the adrenomedullin 1 receptor (AM is the most potent at this receptor) and CLR-RAMP3 produces the AM2 receptor (here AM and AM2 are approximately equipotent). To date the cognate receptor for AM2 and its physiological role remain unknown; no receptor shows marked selectivity for it. While these is an abundance of evidence of GPCR signalling bias in recombinant cell systems, and in this case CLR-mediated bias^12,19^, documented examples using natural agonists and endogenously expressed human receptors are currently lacking. We wished to ascertain whether signalling bias at the CLR occurs in primary cells and whether it plays a role in cellular function. We have chosen to focus our research on RAMP1 and RAMP2 as the CGRP and AM1 receptors are the best described.

Using human endothelial cells which endogenously express the AM1 receptor (CLR-RAMP2), we demonstrate for the first time that biased agonism is present and has a fundamental role in the function of peptide hormones acting on primary human cells. Moreover, through deletion of the endogenous RAMP2 and replacing it with RAMP1, we highlight that not only is the RAMP essential for CLR function and CGRP peptide family signalling in primary cell systems but that RAMPs direct the pattern of agonist bias observed. Furthermore, we document previously unreported actions for the CGRP-based peptide agonists; AM2, in particular, emerges as an agonist uniquely biased to elevate calcium-mediated nitric oxide (NO) signalling while both CGRP and AM display distinct pro-proliferative effects in cardiovascular cells. The work we describe here reveals that GPCR agonist bias occurs naturally in human cells and plays fundamentally important physiological roles.

## Results

### Endothelial cells exclusively express functional CLR/RAMP2 (AM1 receptor)

While there are many reports of biased agonism for GPCRs in recombinant systems (e.g. ^19,20^), few examples have been documented in primary human cells. Given the reported roles of CGRP, AM and AM2 in the cardiovascular system we have focussed our studies upon these peptides, and their receptors in primary human endothelial cells (both HUVECs and human umbilical artery endothelial cells (HUAECs)). Both endothelial cell lines appear to express the AM1 receptor since we could only detect transcripts for CLR and RAMP2 using qRT-PCR (Figure 1 and Figure S1A). This was confirmed functionally, since when the endothelial cells were stimulated with agonists and cAMP accumulation quantified, the rank order of potency was AM>AM2>CGRP (Figure 1B, Figure S1B and Table S1). Furthermore, application of the selective AM1 receptor antagonist AM22-52 which, at 100nM, abolished agonist induced cAMP accumulation while 100nM olcegepant (a CGRP receptor selective antagonist) had little effect (Figure S2A,B). An important factor in confirming receptor-specific agonist bias is to ensure that competing receptors are not present in the system. The closely related calcitonin receptor (CALCR/CTR) not only interacts with RAMPs^21^ but has also be documented to bind CGRP^21^. Endothelial cells appear to not express the CTR since we were unable to detect the presence of its transcript or obtain a significant functional response upon application of two CTR agonists (calcitonin or amylin) (Figure S2C). Thus, based upon these data, we suggested that endothelial cells specifically express the AM1 receptor alone (CLR-RAMP2) and are a useful primary cell line with which to study potential endogenous agonist bias.

**Figure 1.**
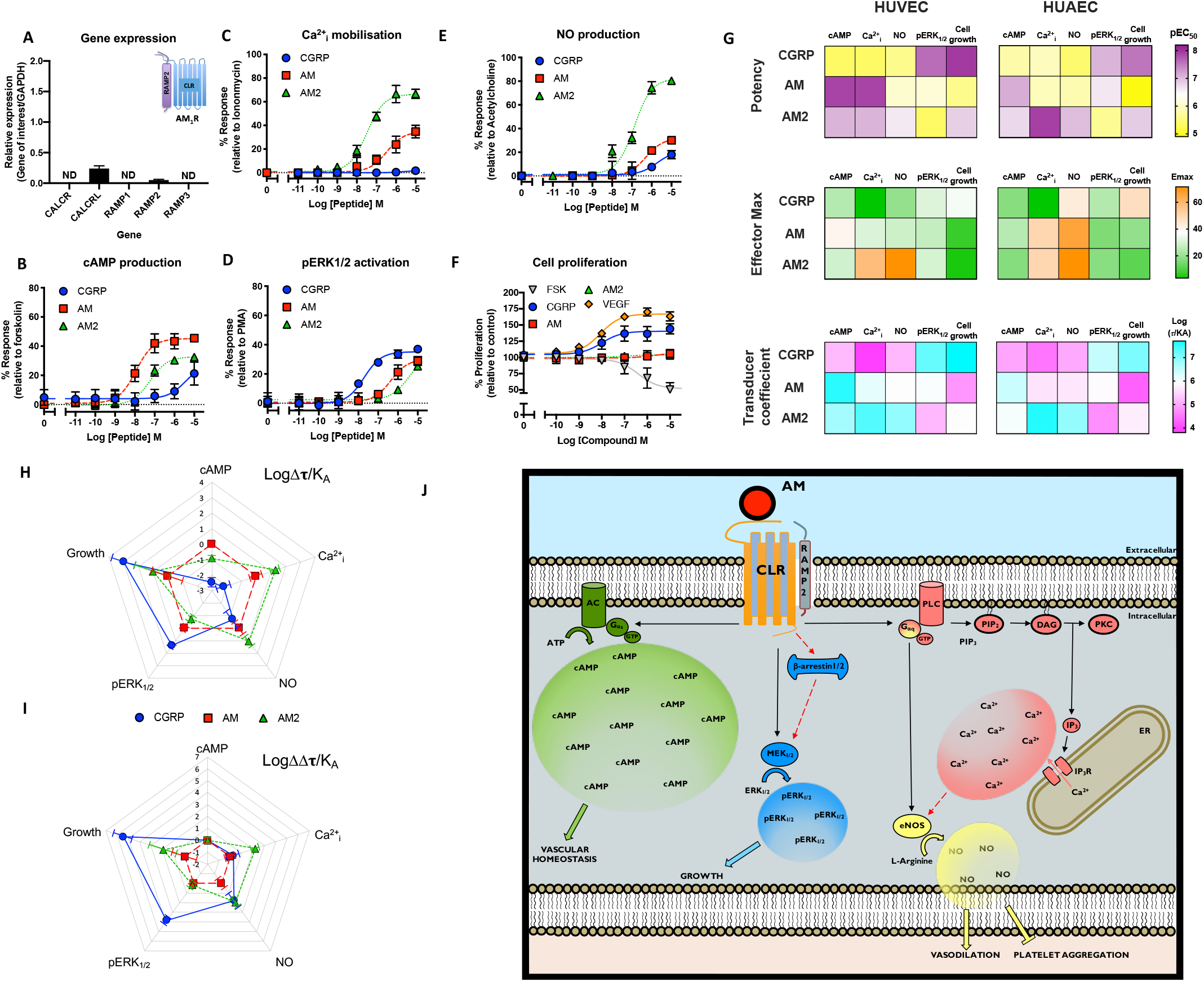
CGRP family peptide signalling bias in HUVECs. **A**) Expression of CALCR, CALCRL, RAMP1, RAMP2, and RAMP3 genes in HUVECs. Data represent mean + SEM of three independent experiments relative to GAPDH expression. ND = not detected in all three samples. **B-F)** Dose-response curves were constructed for HUVECs stimulated with CGRP, AM or AM2 and the cAMP levels quantified relative to forskolin (100μM) (**B**), mobilisation of Ca^2+^_i_ relative to ionomycin (10μM) (**C**), intracellular ERK_1/2_ phosphorylation relative to PMA (10μM) (**D**), total nitric oxide production relative to acetylcholine (10μM) (**E**), and extent of cell proliferation (after 72 hours) relative to vector treated control and VEGF (**F**). Data are analysed using a three-parameter non-linear regression curve or the operational model of receptor agonism^27^. Data are analysed using a three-parameter non-linear regression curve. **G**) Heatmaps representing the signalling properties between HUVEC and HUAEC cells for potency, effector maximum and the transducer coefficient. **H-I**) Signalling bias plots were calculated as ΔLog(τ/K_A_) (**H**) or ΔΔLog(τ/K_A_) (**I**) for each agonist and for each signalling pathway. Determination of values requires normalisation to a reference agonist (AM) alone in **G**, while for **H** values were normalised to both a reference agonist (AM) and a reference pathway (cAMP). All data represent mean ± SEM of at least 3-6 independent experiments. **J**) Representation of the signalling outcomes downstream of CLR-RAMP2 stimulated by adrenomedullin in a HUVEC. Including G proteins known to couple to CLR-RAMP2 and their signalling pathways as a result of AM mediated receptor activation. Solid arrows indicate known pathways. Dashed arrows represent possible pathways.

### Endogenous agonist bias at the CLR-RAMP2 receptor

For studies of biased agonism, it is not simply the ability of different ligands to activate the canonical second messenger pathway to varying extents that is important, but their ability to differentially activate a multitude of downstream pathways. Having established that our primary endothelial cells express only one of the receptor-RAMP complexes responsive to our three peptides: CGRP, AM and AM2, we next sought to quantify the extent of endogenous agonist-induced biased signalling through CLR-RAMP2 at other pathways. Beyond coupling to Gα_s_, in recombinant systems, it has been shown that the CLR can couple to the Gα_q_ family of G proteins to mobilise Ca^2+^_i_ 19. Consistent with these previous reports we were able to observe concentration-dependent increases in Ca^2+^_i_ in both HUVECs and HUAECs upon application of AM and AM2 but little or none with CGRP (Figure 1C, Figure S1C and Table S1). Importantly, all responses could be abolished with the co-treatment of the Gα_q/11/14_ inhibitor YM-254890^22^ (Figure S2D) suggesting the response observed was purely Gα_q/11/14_-mediated and thereby confirming CLR-based pleiotropy in primary endothelial cells. Interestingly, at the CLR-RAMP2 receptor in both endothelial cell lines, AM2 produced the most potent response and not AM suggesting that a non-cognate agonist can have a distinct and more potent effect than the cognate agonist at certain pathways endogenously.

We subsequently turned our attention to the extracellular signal-regulated kinase (ERK) pathway (assayed after 5 minutes stimulation) where we found that, again, the ‘cognate’ agonist (AM) was not the most potent. Perhaps surprisingly, CGRP (the agonist reported to be the least potent at cAMP production at the AM1 receptor) was the most potent at stimulating ERK_1/2_ phosphorylation (Figure 1D, Figure S1D and Table S1). Thus, despite this being designated an AM1 receptor, it is CGRP and not AM that produces physiologically relevant signalling via the ERK_1/2_ pathway.

### Physiological consequences of CGRP-based peptide agonist bias in primary endothelial cells

As we were exploring the AM1 receptor in its native environment, we sought to discover whether the distinct patterns of agonist bias we have observed with CGRP, AM and AM2 were reflected further downstream as physiological bias. Here we considered two potential physiological outcomes with important therapeutic potential – the generation of NO (a vital modulator of vascular homeostasis) and cell proliferation. NO, generated through endothelial nitric oxide synthase (eNOS) in endothelial cells^23^ promotes vasorelaxation/dilation in a cGMP dependent manner^24^. In both HUVECs and HUAECs we observed that all three agonists could evoke NO synthesis in the order of potencies AM2>AM>CGRP (Figure 1E, Figure S1E and Table S1) although both AM and CGRP were partial agonists for this pathway with the potencies closely resembling the trends observed for Ca^2+^_i_ mobilisation. Indeed, a direct correlation between Ca^2+^_i_ mobilisation and NO production in endothelial cells was confirmed through application of YM-254890 which abolished all NO release (Figure S2E). Such observations are consistent with the role of increases of Ca^2+^_i_ concentrations leading to eNOS function^24^ but to the best of our knowledge have not been demonstrated previously for AM2.

Beyond NO production we also measured the long-term cell proliferation (72 hours) response to the three peptides in both endothelial cell lines. Again, the different peptides had varying effects on proliferation, with CGRP most potently promoting cell growth (Figure 1F, Figure S1F and Table S1). This is consistent with the data we describe for phosphorylation of ERK_1/2_ suggesting proliferation is not mediated via a cAMP-dependent pathway. This was further corroborated by the observation that application of the non-selective adenylyl cyclase activator forskolin induced a concentration dependent inhibition of cell proliferation. Together, this data suggests that CLR exerts important cellular effects in a Gα_s_-independent manner thus unveiling previously undocumented abilities for CGRP to promote proliferation in human cells through the AM1 receptor.

To provide a means of comparison of the extent of agonist bias observed in the endothelial cells (Figure 1G), and to remove potential confounding issue of system bias (note, system bias may arise due to the differential expression of signalling components or cofactors in the cellular background of choice) we fitted our data with operational model of receptor agonism^27^ for both endothelial cells (Figure 1H-J and Figure S1G-H and Table S1). There was a strong similarity in the signalling profiles between the two endothelial cells across the five different pathways (Figure 1H) with significant correlations in potency (Figure S3C; *r* = 0.73 – 95% confidence interval 0.35 – 0.90; *p* < 0.01) and the transducer coefficient (Figure 1I) (Figure S3C; τ/K_A_; *r* = 0.94 – 95% confidence interval 0.84 to 0.98; *p* < 0.0001) suggesting primary endothelial cells share common AM1 receptor signalling properties. Finally, this analysis reinforced the notion that AM2 is biased towards Ca^2+^_i_ mobilisation and NO production while CGRP favours pERK_1/2_ activation and cell proliferation.

### AM1 receptor-mediated cAMP accumulation and pERK_1/2_ activation exemplify agonist bias

The mechanism by which adenylyl cyclase is regulated involves competition between G_s_ (activation) and members of the G_i/o_ (inhibition) family of G proteins. Semi-quantitative RT-PCR in both endothelial cell lines revealed the presence of the same Gα subunits (Figure S3D-E) and both β-arrestin1 and 2 in both cell lines including members of G_i/o_ family. We and others have documented how the AM1 receptor (in agreement with other class B GPCRs) can couple to the inhibitory G proteins^21,28,29^ although this is often observed in overexpression systems and is cell type dependent. Application of pertussis toxin (PTX), which ADP-ribosylates the inhibitory G proteins (with the exception of G_z_), to the HUVECs revealed a dose-dependent increase in cAMP accumulation (Figure 2A) and a suppression of ERK_1/2_ phosphorylation upon application of CGRP and AM2 but not AM (Figure 2B). This data, consistent with our previously reported work^19^ suggests that only the non-cognate agonists (CGRP and AM2) are able to recruit G_i/o_ proteins to the CLR, and, particularly in the case of CGRP, the purpose of this is to bias the response away from cAMP and towards other pathways such as pERK_1/2_. This is highly likely to contribute to the differences seen in physiological outcomes such as proliferation, outlined above.

**Figure 2.**
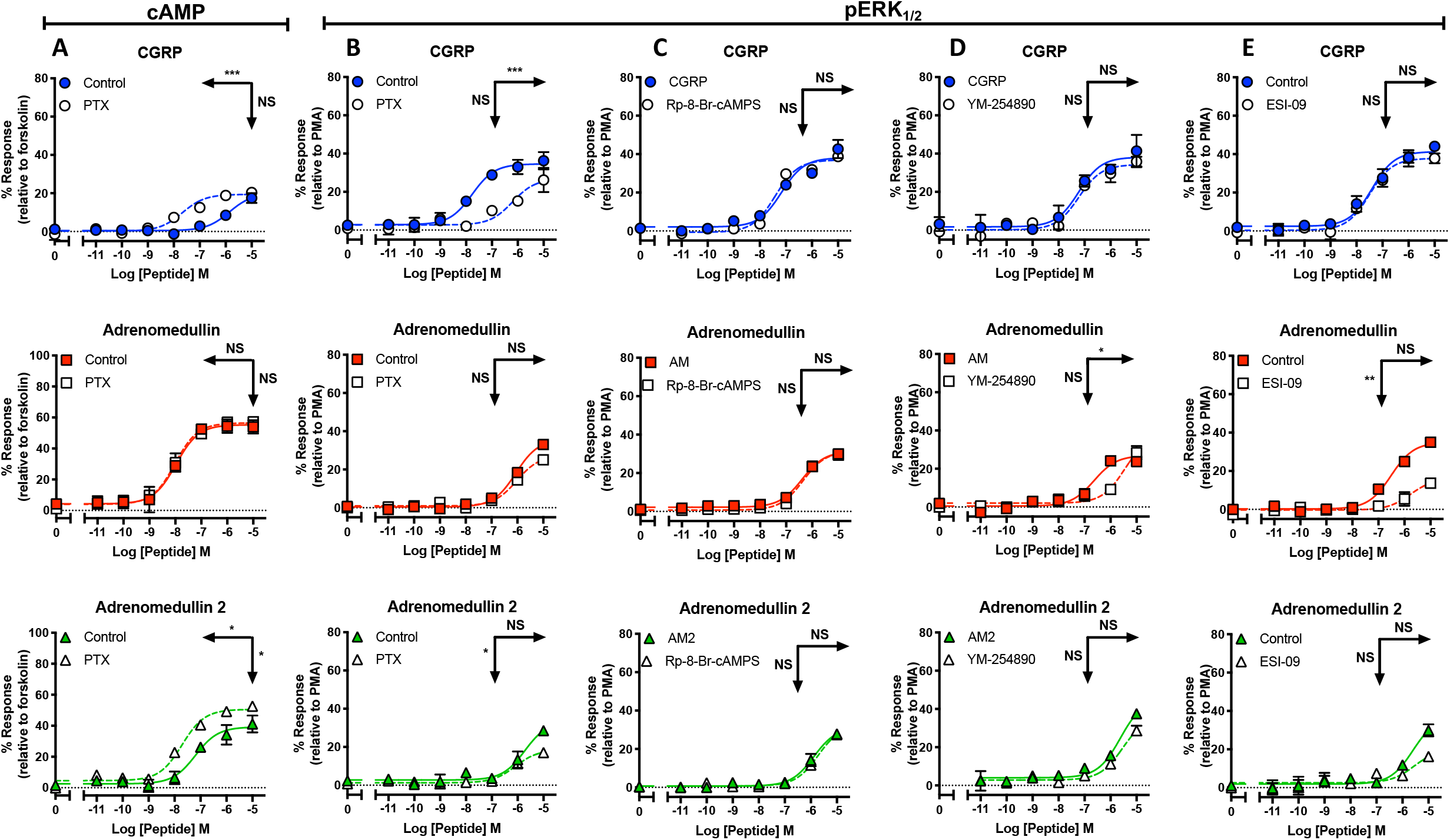
Non-cognate Ga couplings at the CLR-RAMP2 complex modulates cAMP accumulation and ERK_1/2_ phosphorylation. **A**) Characterisation of cAMP accumulation in response to stimulation by CGRP, AM and AM2 with and without PTX treatment relative to forskolin (100μM). **B-E**) Characterisation of ERK_1/2_ phosphorylation in response to stimulation by CGRP, AM and AM2 with and without PTX treatment relative to PMA (10μM) (**B**), with/without YM-254890 relative to PMA (10μM) (**C**), with/without Rp-8-Br-cAMPS (10 μM) (**D**), and with/without ESI-09 (**E**). Data are analysed using a three-parameter non-linear regression curve. All mean values ± SEM are calculated from 3-6 independent experiments. Statistical significance determined compared to control using an unpaired Student’s t test with Welch’s correction (*, p<0.05; **, p<0.01; ***, p<0.001 ****, p<0.0001). NS denotes no statistical significance observed. Rows show pEC_50_, and vertical arrows show E_max_ statistical significance.

This did however pose the question as to how AM modulates the pERK_1/2_ response. Inhibition of protein kinase A (PKA) had no significant effect (Figure 2C) however antagonism of G_q/11/14_ signalling did reduce the potency of AM-mediated pERK_1/2_ activation (Figure 2D). More strikingly, inhibition of the exchange proteins directly activated by cAMP (EPAC)1/2 activation significantly attenuated both the potency and magnitude of the maximal response (Figure 2E). Taken together, these data highlight the wide array of different G protein couplings and their interlinking actions upon downstream signalling events for the AM1 receptor. These couplings have not been engineered so are not enhanced by overexpression artefacts and thereby represent pure endogenous agonist bias.

### RAMP isoform is essential for CLR-mediated agonist bias

One of the advantages of using recombinant cell lines and/or model cell organisms is the ability to switch the expressed GPCR or RAMP to observe effects on agonist bias. However, these recombinant systems do not allow for observations of physiological bias. Thus, we next sought to determine the effects of CGRP-based agonist bias in primary cells where the endogenous RAMP had been switched using gene deletion followed by lentiviral reintroduction. We chose the HUVEC cell line with which to perform the gene editing since we have been able to grow HUVECs beyond passage P6 to P14 before loss of CGRP based signalling responses are observed (Figure S4), and it is necessary to grow them past P6 to develop the RAMP2 null cells. We used lentiviral CRISPR-Cas9 to knockout the RAMP2 gene from the HUVECs. We used a pooled sgRNA strategy using three sgRNAs in separate lentivirus (Figure S5A) which were selected using a puromycin resistance cassette (Figure S5B) to increase our efficiency of editing. We transduced HUVECs at a high multiplicity of infection (MOI) of 10, ensuring that each cell was infected by several lentivirus to increase the likelihood of achieving a deletion. Then sgRNA editing efficiency in the remaining cell pool was assessed by PCR amplification of targeted region, Sanger sequencing and TIDE^30^ analysis: all demonstrating an editing efficiency greater than 95%. This was then followed by analysis by qRT-PCR for receptor and RAMP mRNA. The expression of CLR remained, but RAMP2 was lost suggesting degradation through nonsense-mediated mRNA decay (NMD) (Figure S5C). It is also important to note that the gene editing did not have any impact upon the cells rate of proliferation (Figure S5E) and the Ga subunit/β-arrestin profile also remained consistent with the wild type HUVECs (Figure S5D). Finally, convinced that the cells were as near to wild type HUVECs as possible, except lacking RAMP2, we assessed their signalling properties following stimulation with all three agonists. In all cases the responses were abolished, although the extent of signalling for the positive controls remained intact (Figure S5F-J). Thus, these data confirm, that loss of RAMP2 in HUVECs abolishes CLR function.

We next introduced, using lentiviral overexpression and blasticidin selection, the open reading frame of RAMP1 into our HUVECΔRAMP2 cell line so, in effect, switching the expressed GPCR from the AM1 receptor to the CGRP receptor. mRNA levels were quantified demonstrating successful introduction of a high level of RAMP1 expression (Figure 3A). We next performed cAMP accumulation assays using CGRP, AM and AM2 confirming that a functional CLR receptor was formed in these modified HUVECs (Figure 3B and Table S2). Reassuringly, we now observed that CGRP was the most potent agonist for the stimulation of cAMP – as expected for a cell line expressing the CGRP receptor (CLR-RAMP1). Perhaps more interestingly, CGRP was also the most potent at mobilising Ca^2+^_i_ (Figure 3C and Table S2) and this was also the case in the associated NO production (Figure 3E and Table S2). Comparison of the ERK_1/2_ phosphorylation (Figure 3D and Table S2) highlighted that AM was now the most potent agonist; a clear switch from wild type HUVEC cells where CGRP was the most potent. This followed to proliferation where AM was also the most potent ligand, although both AM2 and CGRP could also promote growth (Figure 3F and Table S2) and was in contrast to the wild type HUVECs where neither could cause proliferation. Thus switching the RAMP in the HUVEC cell line appears to have had a dramatic effect on the agonist bias observed and the functional consequence (Figure 3G,H and Table S2) – beyond just cAMP accumulation as would be excepted. However, it should be noted that as RAMP1 expression was high we should be cautious in our direct comparisons between the wild type HUVECs and our RAMP1-HUVEC cell line.

**Figure 3.**
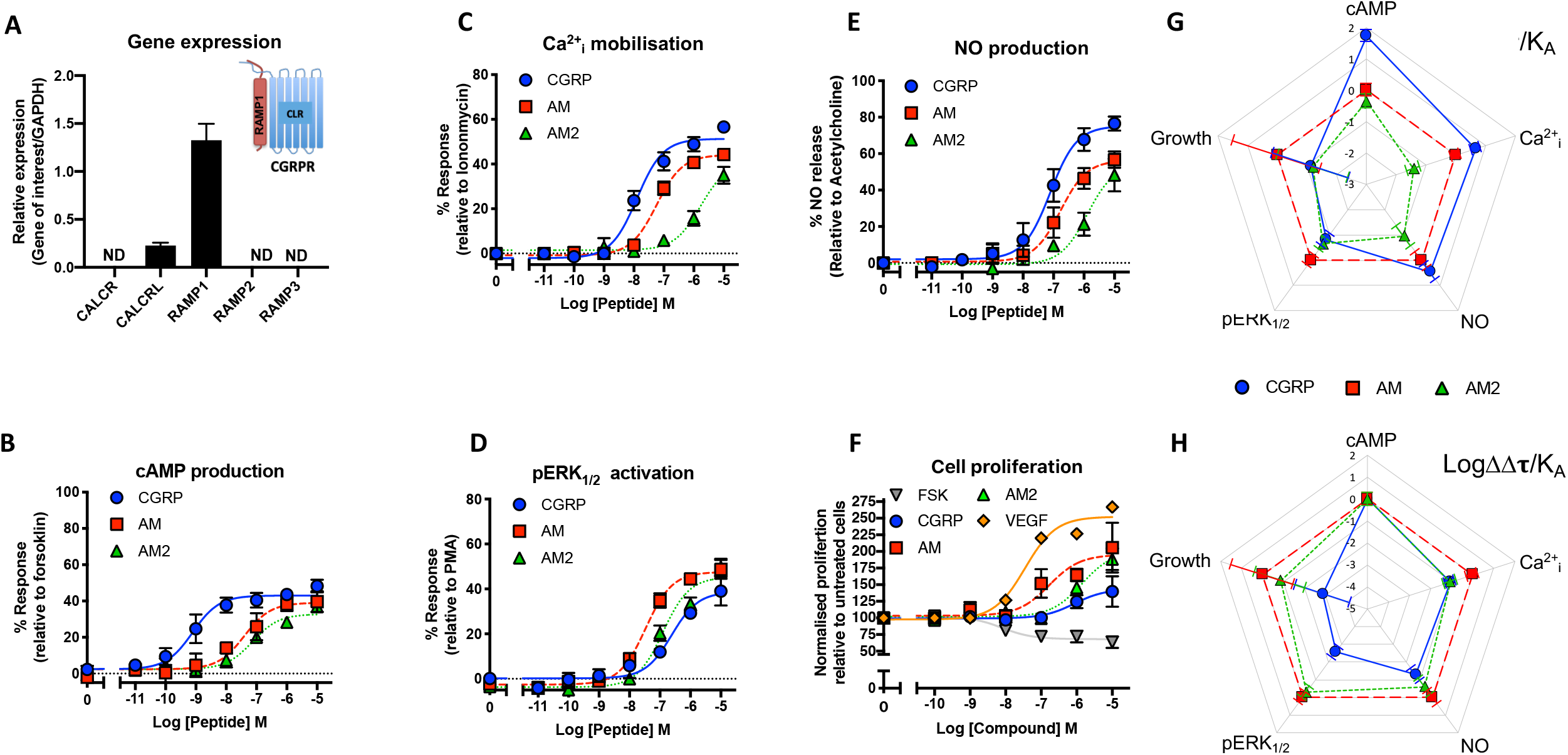
Switching RAMP expression in HUVECs produces unique signalling bias patterns for CGRP family of agonists. **A)** Expression of CALCR, CALCRL, RAMP1, RAMP2, and RAMP3 genes in RAMP1 expressing HUVECs. Data represent mean + SEM of at least three independent experiments relative to GAPDH expression. ND = not detected in all three samples. **B-F**) Dose-response curves were constructed for RAMP1 expressing HUVECs stimulated with CGRP, AM or AM2 and the cAMP levels quantified relative to forskolin (100μM) (**B**), mobilisation of Ca^2+^_i_ relative to ionomycin (10μM) (**C**), intracellular ERK_1/2_ phosphorylation relative to PMA (10μM) (**D**), total nitric oxide production relative to acetylcholine (10μM) (**E**), and extent of cell proliferation (after 72 hours) relative to vector treated control and VEGF (**F**) Data are analysed using a three-parameter non-linear regression curve or the operational model of receptor agonism^27^. Data are analysed using a three-parameter non-linear regression curve. **G-H**) Signalling bias plots were calculated as ΔLog(τ/K_A_) (**G**) or ΔΔLog(τ/K_A_) (**H**) for each agonist and for each signalling pathway. Determination of values requires normalisation to a reference agonist (AM) alone in **G**, while for **H** values were normalised to both a reference agonist (AM) and a reference pathway (cAMP). All data represent mean ± SEM of at least 3-6 independent experiments.

### Endogenous agonist bias at the CLR-RAMP1 in primary human cardiac myocytes

In order to provide a comparison for RAMP1-HUVEC signalling with a primary cell line that endogenously expressed the CGPR receptor we turned to primary human cardiomyocytes (HCMs) since these cells only expressed CLR and RAMP1 (Figure 4A); analogous to endothelial cells, HCMs also do not express a functional CTR (Figure 4A and Figure S6A). To confirm that the mRNA expression translated to functional receptor expression we performed cAMP accumulation assays for the CGRP family of peptides (Figure 4B and Table S2). Here, CGRP was the most potent agonist followed by AM2 and AM, a pattern consistent with the expression of the CGRP receptor^20^ (also confirmed by application of 100nM olcegepant to inhibit cAMP accumulation for all three agonists (Figure S6B) while 100nM AM22-52 (Figure S6C) had little effect). Intriguingly, upon application of PTX to HCMs we were unable to observe any significant change in the potency or maximal signalling for any of the three peptide agonists (Figure S6D) although the transcript for Gα_i2_ was lower than in the endothelial cells (Figure S5E) and this Gα subtype has previously been suggested to be important for PTX-sensitive effects from CLR^19^. In contrast to the wild type HUVECs but analogous to the RAMP1-HUVEC cells, not only was CGRP able to stimulate G_q/11/14_-mediated-Ca^2+^_i_ mobilisation in HCMs but it was the most potent agonist (Figure 4C and Table S2). When quantifying ERK_1/2_ phosphorylation we again observed that the cognate ligand (CGRP) was not the most potent (Figure 4D and Table S2), but as in HUVECs, it was the least potent ligand at cAMP accumulation that was the most potent for ERK_1/2_ phosphorylation. AM was the most potent at stimulating ERK_1/2_ phosphorylation, demonstrating that it can produce functionally relevant signalling responses at the CLR-RAMP1. The order of potency for the three agonists for ERK_1/2_ phosphorylation was replicated in the long-term cell proliferation assays (Figure 4F and Table S2) with AM remaining the most potent. All three peptide agonists could also evoke G_q/11/14_-mediated-NO production (Figure 4E, Figure S6F,G and Table S2) although their responses were less distinct from each other.

**Figure 4.**
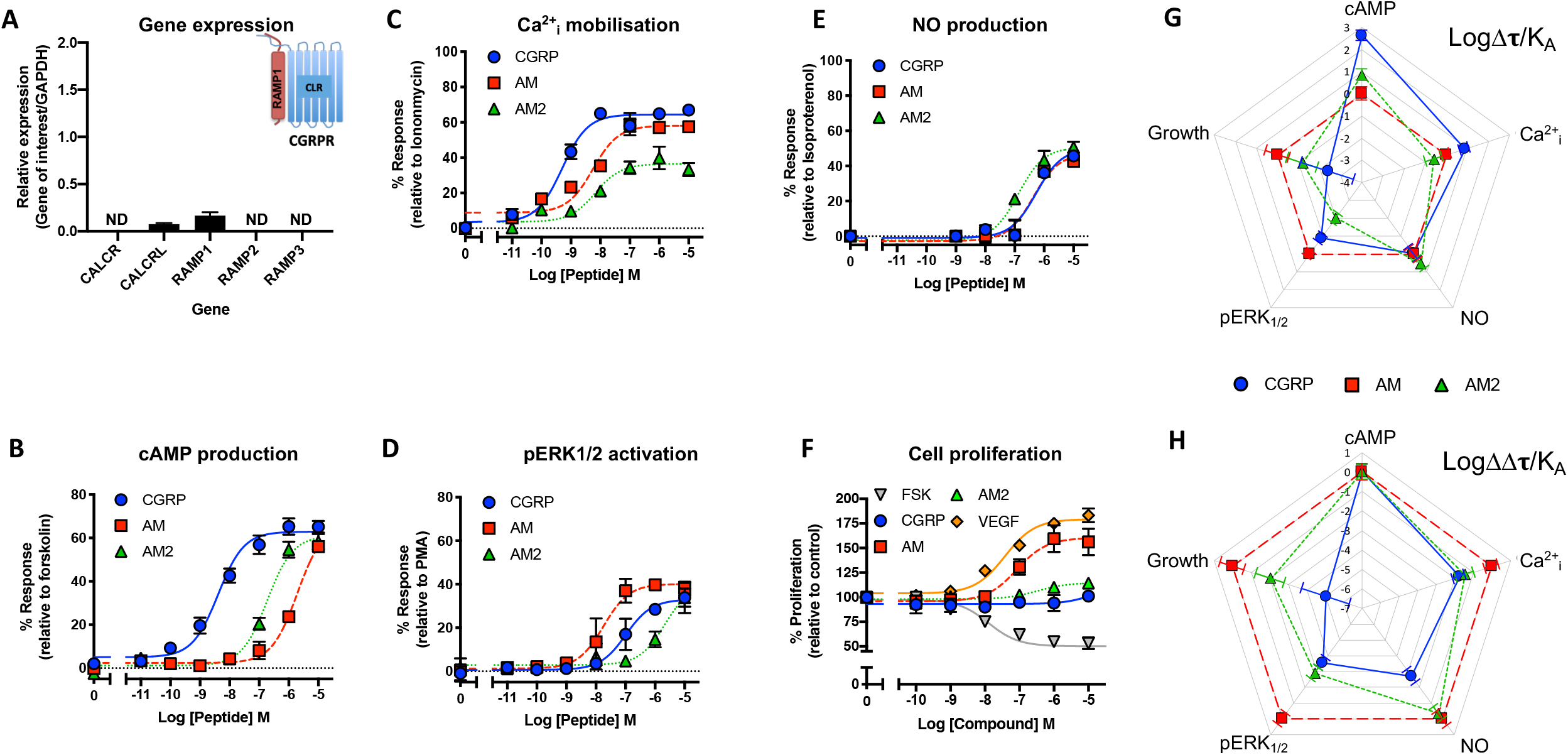
CGRP family peptide signalling bias in human cardiomyocytes. **A)** Expression of CALCR, CALCRL, RAMP1, RAMP2, and RAMP3 genes in HCMs. Data represent mean + SEM of three independent experiments relative to GAPDH expression. ND = not detected in all three samples. **B-F)** Dose-response curves were constructed for HCMs stimulated with CGRP, AM or AM2 and the cAMP levels quantified relative to forskolin (100μM) (**B**), mobilisation of Ca^2+^_i_ relative to ionomycin (10μM) (**C**), intracellular ERK_1/2_ phosphorylation relative to PMA (10μM) (**D**), total nitric oxide production relative to acetylcholine (10μM) (**E**) and extent of cell proliferation (after 72 hours) relative to vector treated control and VEGF (**F**) Data are analysed using a three-parameter non-linear regression curve or the operational model of receptor agonism^27^. Data are analysed using a three-parameter non-linear regression curve. **G-H**) Signalling bias plots were calculated as ΔLog(τ/K_A_) (**G**) or ΔΔLog(τ/K_A_) (**H**) for each agonist and for each signalling pathway. Determination of values requires normalisation to a reference agonist (AM) alone in **G**, while for **H** values were normalised to both a reference agonist (AM) and a reference pathway (cAMP). All data represent mean ± SEM of at least 3-6 independent experiments.

Analysis of the RAMP1-HUVEC signalling profile suggested a close overlap with the properties of HCMs (Figure 5 and Table S2) for the five different signalling pathways as well as an opposing signalling profile to HUVECs. We confirmed this using correlation plots (Figure 5A-F) of both potency and the transducer coefficient Log(τ/K_A_), obtained from application of the operational model of receptor agonism^27^ (values can be found in Table S2 and Table S3). Whilst a positive correlation was detected between RAMP1-HUVECs and HCMs (potency; *r* = 0.55 – 95% confidence interval, 0.051 to 0.82; *p* < 0.05; transducer coefficient; *r* = 0.52 – 95% confidence interval, 0.009 to 0.81; *p* < 0.05), a negative correlation was observed between HUVECs and RAMP1-HUVECs (potency; *r* = −0.54 – 95% confidence interval, −0.83 to −0.04; *p* < 0.05; transducer coefficient; *r* = −0.58 – 95% confidence interval, −0.84 to −0.10; *p* < 0.05). We did not observe any correlations between HUVECs and HCMs which, in part appears due to the HCMs having reduced capacity to release NO. Finally, when we extended our analysis to determine the change in transducer coefficient normalised to cAMP accumulation mediated by the non-cognate (for RAMP1-CLR or RAMP2-CLR) agonist AM2 (Figure 5G and 5H) it becomes apparent how closely aligned the signalling properties in RAMP1-HUVECs and HCMs are.

**Figure 5.**
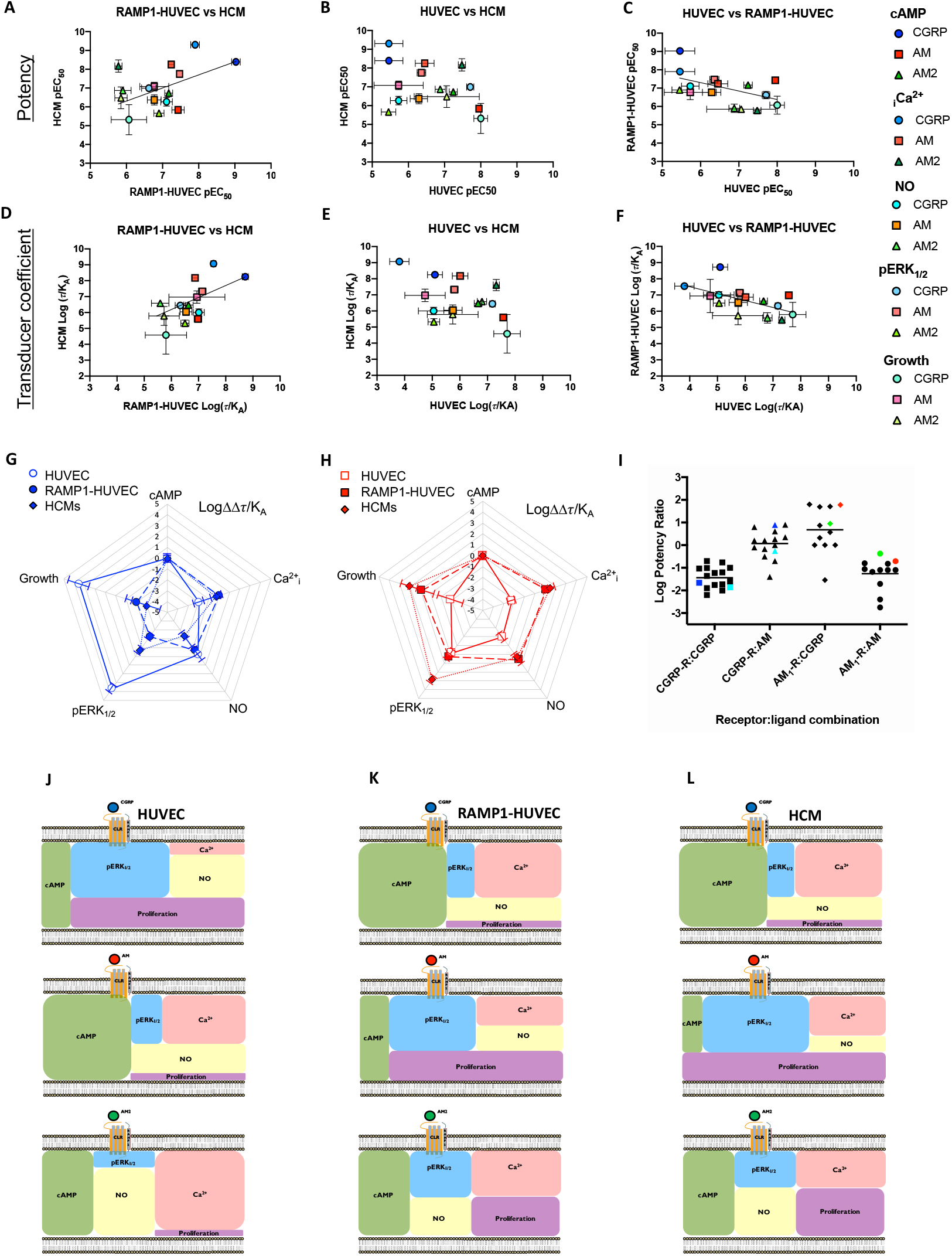
CGRP signalling bias in RAMP1 expressing HUVECs correlates with that in human cardiomyocytes. **A-C**) the correlation of Log agonist potencies ± SEM for CGRP, AM and AM2 stimulated cAMP accumulation, mobilisation of Ca^2+^_i_, NO production, intracellular ERK_1/2_ phosphorylation and cell proliferation in RAMP1 expressing HUVECs and HCMs (**A**), HUVECs and HCMs (**B**) and HUVECs and RAMP1 expressing HUVECs (**C**) was analysed by a scatter plot and Pearson’s correlation coefficients (*r*) were calculated. A significant positive correlation was observed for RAMP1 expressing HUVECs and HCMs. The presence of a line indicates a positive correlation. **D-F**) as for **A-C** except the transduction coefficient Log (τ/K_A_) was calculated. A significant positive correlation was observed for RAMP1 expressing HUVECs and HCMs as indicated by the presence of a line. **G)** Signalling bias plots were calculated as ΔΔLog(τ/K_A_) for CGRP in the three cell lines, HUVECs, RAMP1 expressing HUVECs and HCMs for each pathway. Values have been normalised to a reference agonist (AM2) and the reference pathway (cAMP) for all three cell lines. **H**) as for **G** except the calculated values are for AM. **I**) Log potency ratios (as measured by the accumulation of cAMP) calculated as Log (EC_50_AM2/EC_50_ agonist). Data are compiled from^12,19^. HUVECs and HUAECs are shown in red and green respectively, HCMs in cyan and RAMP1-HUVECs in blue. **J-L**) Schematic representation of the signalling bias produced by CGPR (**J**), AM (**K**) and AM2 (**L**), and the intracellular ‘signalling codes’ they bring about based on the potencies recorded at individual pathways in HUVECs, RAMP1-HUVECs and HCMs.

## Discussion

We have shown for the first-time that the CGRP family of endogenous peptides demonstrate biased agonism at the endogenous CLR in a physiological system; and that the RAMP expressed dictates the intracellular response and ultimately the physiological outcome (Figure 5). Many receptors have been shown to demonstrate agonist bias; but for the most part this has been shown through synthetic ligands designed to target certain receptor pathways^2^. We have now shown that this is a process that can occur physiologically to direct different outcomes. Through elucidating distinct patterns of signalling bias that each peptide-receptor-RAMP produces, we have shown that bias is a naturally occurring phenomenon in a range of human cardiovascular cells. Furthermore, while we have only begun to scratch the surface of how important bias is physiologically, it is now clear it is an intrinsic part of endogenous CLR function, and we anticipate this is the case for many more GPCRs that exhibit signalling bias in over-expression studies. We have also demonstrated the importance of studying GPCR second messenger signalling with the endogenous receptor in its native environment, with the distinct signalling patterns of AM2 we have uncovered providing a good example of this.

We have confirmed, as anticipated by the co-expression models, that endogenous CLR is unable to function without RAMP expression through CRISPR-Cas9 KO of the endogenous RAMP. This provides, to the best of our knowledge, the first example of CRISPR-Cas9 interrogation of GPCR function in a primary cardiovascular cell. Furthermore, we have shown that the expression of a different RAMP in the HUVECs can switch the signalling bias of the CLR and associated peptide agonists, thus providing additional evidence that RAMP targeting could become a powerful therapeutic tool^25^.

We have compared the pharmacology of these receptors in terms of cAMP accumulation to reports compiling multiple values from independent publications using the human receptor in transfected systems^20^. It was reassuring therefore to observe that CGRP, AM, and AM2 displayed similar trends in cAMP potency at CLR-RAMP1/2 in the primary cells to those seen in recombinant co-expression studies, as well as in the gene edited RAMP1-HUVECs which reflected the cAMP data from transfected systems (Figure 5I). In addition, we have performed a comprehensive analysis of the mechanisms used by RAMP2-CLR complexes to stimulate ERK_1/2_ phosphorylation in endothelial cells (Figure 2). It is apparent that each agonist uses unique mechanisms to activate ERK_1/2_. For both CGRP and AM2 it is mediated in a G_i/o_-dependent manner, while AM uses a combination of G_q/11/14_ signalling and EPAC activation. None of the agonists appear to mediate their ERK_1/2_ stimulation through the so-called cognate pathway, cAMP accumulation. The data presented is ERK_1/2_ phosphorylation after 5 minutes and it will be of interest to determine the mechanisms and spatial locations that facilitating long term ERK_1/2_ phosphorylation.

What has become clear in this study is that each of the endogenous ligands have very specific potencies at each pathway measured, whether it is at CLR-RAMP2 in HUVECs/HUAECs or CLR-RAMP1 in HCMs or RAMP1-HUVEC cells. Each peptide generates their own unique signalling profile (Figure 5J-L). In a concentration-dependent manner, they individually recruit distinct G proteins in a manner regulated by the RAMP. This leads to a specific pattern of second messenger production and therefore a ‘signalling barcode’ for the cell to interpret and produce further physiologically necessary downstream responses. Expression analysis reveals in our three primary cell lines that each only expresses mRNA above the detection threshold for one RAMP and the CLR. Combined with the cAMP signalling profile for each it appears that endothelial cells and HCM are a excellent primary model cell types for the analysis of how the CLR-RAMP2 and CLR-RAMP1 signal *in vivo*.

It is worth noting that the present method of classifying receptors for CGRP and AM is based upon their potencies at cAMP production in addition to their affinities in binding assays^26^. This method arose due to the assumption that cAMP was the most physiologically relevant pathway. Here, we have demonstrated that significantly different potencies are observed for agonists and these lead to physiologically relevant outcomes. As such we need to carefully consider how we classify CGRP-related receptors in the future, and more widely all GPCRs that exhibit agonist bias.

We can also consider this work in the wider context of the organs and systems these cells are found in as this sheds light on some of the pathways, and involvement of bias in some of the established roles of CGRP family peptides in the cardiovasculature. It has long been recognised that all three peptides show pleiotropic signalling, activating multiple G proteins and signalling pathways^33^ (and indeed this continues in current literature^34-36^), but it has previously not been possible to fit this into any framework. We suggest our current observations on RAMP-directed bias may assist with this.

AM has a multitude of important roles in vascular homeostasis^6^; one of which is regulating endothelial barrier function^37^. It is thought to cause barrier stabilisation and protect against infection mediated junctional protein disappearance, all brought about initially through cAMP production^38^. This is supported by our work demonstrating AM produces a potent cAMP response and is biased towards this pathway. It is also well documented that AM is a potent vasodilator known to mediate some of its vasodilatory effects through NO release from vascular endothelial cells^25,39^ which we have pharmacologically profiled here.

In contrast, the precise role of AM2, which is also found in endothelial cells, has been unclear. We now provide evidence that AM2 is a potent stimulator of Ca^2+^_i_ mobilisation and NO synthesis. It is possible therefore that this plays a vital role at least in umbilical endothelial cell physiology, and indeed wider vascular physiology. Thus, the novel finding of AM2’s greater potency than AM at eliciting NO release via Ca^2+^_i_ mobilisation may have great therapeutic potential.

Interestingly, in vascular endothelial cells CGPR inhibits adenylyl cyclase through G_i/o_ and predominantly signals through pERK_1/2_, and proliferation. The link between pERK_1/2_ and cellular metabolism/proliferation is well established^40-42^, as well as in endothelial cells specifically^43-45^. We have shown that where an agonist has biased signalling towards ERK_1/2_ phosphorylation, this is carried through to long term cellular proliferation. Importantly, this shows overall that two non-cognate ligands, often not considered significant for receptor function, do in fact have important signalling and physiological roles/capabilities. In addition, we have demonstrated that endogenous pERK_1/2_ can come from a variety of sources depending on the stimulating ligand. Together this shows that the CLR initiates a multitude of intracellular pathways beyond simply G_s_ and cAMP/PKA in physiologically relevant cells. Therefore, our data adds further evidence that AM for CLR-RAMP2 and CGPR for CLR-RAMP1 should only be considered the cognate ligands in terms of G_s_-mediated cAMP signalling, and when looking at the physiology of RAMP2 in endothelial cells and the vasculature as a whole, CGRP and AM2 should be considered alongside AM for their different and potentially complementary roles.

On the heart, there are multiple reports that CGRP has a cAMP-mediated positive inotropic and chronotropic effect^5,46,47^, and our data showing its strong response in cAMP accumulation assays on human myocytes (combined with its overall bias towards this pathway) supports this. There are contrasting reports in the literature over AM’s effect on heart contractility^7,48^; with some suggestions that it is a positive inotrope acting in the same cAMP driven manner as β-adrenoceptor agonists^49^, while others report it having negative inotropic effects^50,51^. Here we show that AM promotes a cAMP response through CLR-RAMP1 in human cardiomyocytes, but it has weak potency. This may provide some context/explanation for the contradictory literature reports. Furthermore, our report has clearly revealed that cAMP is not AM’s primary signalling pathway in HCMs and that it is biased towards pERK_1/2_ and cell proliferation rather than cAMP and positive inotropy. Nevertheless, evidence suggests that AM has an important role in the human heart. This includes the observed elevation of AM in the failing heart^52^. Here, we have utilised proliferating human ventricular myocytes *in vitro* and shown that AM (but not CGRP or AM2) exhibits signalling bias specifically towards pERK_1/2_ and enhancing proliferation in these cells. This work highlights AM as a novel peptide hormone that may promote cardiac regeneration naturally *in vivo*, and provides a cellular mechanism for this. This may also explain the elevation of AM in heart failure^6^ and the clinical trial data showing that AM administration reduces infarct size^27^. For AM2, its effect on contraction of the heart is blocked by both inhibitors of PKA and PKC (i.e. G_q/11/14_-coupling), suggesting multiple signalling pathways are activated by this peptide *in-vivo*^54^.

It should be noted that not all studies reveal pleiotropic signalling for the CGRP family of peptides, either involving direct measurements of G protein coupling in cell-free systems^55^ or second messenger generation in cells^12^. We suggest that there is cell membrane (e.g. lipid composition) and cell line-specific factors (e.g. expression of G proteins) that influence the observed bias.

In summary, we have gone beyond previous studies in recombinant systems to observe agonist bias. While we have focused upon CLR-RAMP complexes, our ability to switch the RAMP means we are, in effect, switching the expressed receptor and therefore has general applicability to all GPCRs. Our data highlights how endogenous agonist bias can have profound consequences for the cell and how important cell background is in regulating this process. This work may even go as far as to suggest that to fully understand bias at a GPCR, it has to be considered in its native environment. While our work takes an important step closer to understanding how the CGRP family of peptides and receptors function on a cellular level in the human cardiovascular system, it also highlights the importance of endogenous agonist bias as a concept and emphasises its long-term consequences for drug design.

## Materials and Methods

### Cell Culture

HUVECs and HUAECs (were both sourced from PromoCell, Germany) were cultured in Endothelial Cell Growth Media (ECGM) (PromoCell). Human Cardiac Myocytes (PromoCell,) were grown in Cardiac Myocyte Growth Media (CMGM) (PromoCell). All cell lines were cultured in media containing 10% heat inactivated Fetal Bovine Serum (FBS) (Sigma, USA). Cells were grown in 25cm^2^ flasks or 75cm^2^ flasks depending on cell density required. They were passaged approximately every 4 days depending on confluency with a final volume of 10ml produced from 1ml of the previous cell culture and 9ml of the growth medium in 75cm^2^ flasks, or 1ml using 4ml in 25cm^2^ flasks, and used from passage 2-6 (with the exception of HUVECΔRAMP2 and RAMP1-HUVEC). All cells grown with 1% antibiotic antimycotic solution (100x Sigma, USA). The cells were maintained in an incubator (37 °C, humidified 95% air, 5% CO_2_) between passaging.

### Genome Engineering

HUVECs with the RAMP2 gene knocked out were generated by CRISPR/Cas9 homology directed repair as described previously^56^. The sgRNA sequences were designed (5’-CGCTCCGGGTGGAGCGCGCCGG-3’), (5’-TCCGGGTGGAGCGCGCCGGCGG-3’), and (5’-CCCGCGTCTCCCTAGGACCCGA-3’) for Cas9 targeting to the human RAMP2 gene (Sigma, US). All guides were delivered in the LV01 vector (U6-gRNA:ef1a-puro-2A-Cas9-2A-tGFP) vector provided by (Sigma-Aldrich, US). Sequences were verified by Sanger sequencing. The control cell line was established by transduction of LV01 vector not containing sgRNA targeted to RAMP2 gene. HUVEC cells were seeded in 6 well plates at a cell density of 160,000 cells/well and maintained at 37°C in 5% CO_2_ with Complete Endothelial Cell Growth Media containing 100μg/ml streptomycin (Sigma-Aldrich, US). 24hrs after seeding virus containing individual sgRNA/Cas9 constructs were pooled and transduced into cells at a high MOI of 10, ensuring that each cell is infected by several lentivirus and increasing the likelihood of achieving KO. Transduction was performed in media containing 8μg/ml Polybrene (Sigma, USA). Cells were cultured for 24hrs then treated with Puromycin (1μg/ml) (Thermo Fisher Scientific, UK) for 3 days to select for transduced cells. Cells then cultured without puromycin and expanded before cells were collected for genotyping by Sanger sequencing, qRT-PCR, and functional assays. All data shown were from cells expanded from these colonies. RAMP1 expression achieved through transduction of virus containing RAMP1 MISSION TRC3 Open Reading Frame (ORF) plasmid (pLX_304) (Sigma, US) into RAMP2 KO-HUVECs. HUVEC cells were seeded in 6 well plates at a cell density of 160,000 cells/well and maintained at 37°C in 5% CO2 with Complete Endothelial Cell Growth Media containing 100μg/ml streptomycin (Sigma, US). 24hrs after seeding, virus containing the ORF construct was transduced into cells in media containing 8μg/ml Polybrene. Cells were cultured for 24hrs then treated with blasticidin (5μg/ml) (Thermo Fisher Scientific, UK) for 6 days to select for transduced cells. Cells were collected for genotyping by qRT-PCR and expanded for functional assays. All ‘HUVEC RAMP1’ data shown were from cells expanded from these colonies.

### Immunofluorescence

HUVEC cells were seeded in Cell Carrier Ultra 96 well plate (Perkinelmer, Boston, MA, US) at a cell density of 160,000 cells/well and maintained at 37°C in 5% CO_2_ with Complete Endothelial Cell Growth Media containing 100μg/ml streptomycin. Cells were washed twice with PBS, fixed with 4% paraformaldehyde (PFA) in PBS (10 mins, RT) then washed three times with PBS. The cells were permeabilized with 0.05% Tween 20 in PBS (60 mins, RT), and then incubated with 10% goat serum in PBS (60mins, RT). The cells were then incubated in primary antibody for cas9 protein (Cell signalling technology, MA, US) 7A9-3A3, diluted 1/700 in PBS/0.05% Tween/3% BSA) at 4°C, overnight and protected from light. The cells were washed three times with PBS and incubated with AlexaFluor 488 goat anti-mouse (Invitrogen A11001, 1/500) (1hr, RT) protected from light. Cells were washed three times with PBS, then nuclei were stained with Hoechst (Invitrogen) (1/2000 in PBS, 10mins, RT). Cells were then washed three times with PBS w/o Mg^2+^ or Ca^2+^_i_ and imaged at 20x magnification (Cell Voyager 7000S, Yokogawa).

### Sequencing of genomic loci

Genomic DNA was extracted from virally transduced HUVEC cells by: collecting approximately 10,000 cells, washing in PBS (sigma-Aldrich, US) and then lysing with DirectPCR Lysis Reagent (Viagen Biotech, US) containing Proteinase K (Qiagen, Germany) at 0.4mg/ml. The lysate was incubated at 55°C for 4 hrs; 85°C, for 10mins; 12°C for 12hrs. PCR reaction was then set up in (20μl) as follows: 2x Flash Phusion PCR Master Mix (Thermo Fisher, US) (20μl), forward primer (5’-AATTCGGGGAGCGATCCTG −3’) (Eurogentec, Belgium) (1μl)(10μm), reverse primer (5’-GAGACCCTCCGAAAATAGGC −3’) (Eurogentec, Belgium) (1μl)(10μm), DNA (100ng/μl)(1μl), ddH2O (7μl). The product was amplified by PCR using the following program: 98°C, 1min; 35x (98°C, 10secs; 55°C, 10secs; 72°C, 15secs), 72°C, 1min; 4°C, hold. PCR clean-up was performed prior to sequencing using the Illustra GFX PCR DNA and Gel-band Purification Kit (Illustra, Germany) according to manufacturer’s instructions. Editing of RAMP2 gene was confirmed by Sanger sequencing (Eurofins) and TIDE analysis^30^.

### Quantitative real-time reverse transcription polymerase chain reaction (qRT-PCR)

HUVECs were cultured as above in Complete Endothelial Cell Growth medium and plated in a 24 well plate at 100,000 cells/well. Media was then removed, and cells were washed in PBS (Sigma, UK). RNA was extracted and genomic DNA eliminated using an RNA extraction kit (QIAGEN, Germany) as per manufacturer’s instructions. The yield and quality of RNA was assessed by measuring absorbance at 260 and 280 nm (Nanodrop ND-1000 Spectrophotometer, NanoDrop technologies LLC, Wilmington DE USA). RNA was used immediately for the preparation of cDNA using the Multiscribe reverse transcriptase. For the preparation of cDNA 100ng of RNA was reverse transcribed using Taq-man reverse transcription kit (Life Technology, MA, USA) according to manufacturer’s instructions. Reactions were performed on a thermal Cycler as following: 25°C, 10mins; 48°C, 30mins; 95°C, 5mins. cDNA was stored at −20°C.

For each independent sample, qPCR was performed using TaqMan Gene Expression assays according to manufacturer’s instructions (Life Technologies, MA, USA) for GAPDH (Hs02786624_g1), CALCR (Hs01016882_m1), CALCRL (Hs00907738_m1), RAMP1 (Hs00195288_m1), RAMP2 (Hs01006937_g1), RAMP3 (Hs00389131_m1) and plated onto fast microAmp plates containing 2μl cDNA, 1μl Taq-man probe, 10μl Taq-man fast universal master mix (Applied Biosystems) and 10μl ddH2O. PCR reactions were performed on ABI 7900 HT real time PCR system (Thermo Fisher Scientific, UK). The program involved the following stages: 50°C, 2 mins; 95 °C, 10mins, the fluorescence detection over the course of 40x (95°C, 15secs; 60°C, 1min). Data are expressed as relative expression of the gene of interest to the reference gene GAPDH where: Relative expression = 2-((Cq of gene of interest) – (Cq of GAPDH)). For the genes where no mRNA was detected these samples are omitted from graphs and labelled accordingly, as indicated in the figure legends.

### In vivo assays

#### Measurement of intracellular cAMP

All primary cell lines were cultured as above. On the day of the experiment media was removed and cells washed with PBS, before being dissociated with Trypsin-EDTA 0.05% (Gibco, UK) and then resuspended in PBS/BSA (0.1%) (Sigma, UK). Cells were immediately plated for use in cAMP assay as per manufacturer’s instructions and as described previously^21^, reagents used were provided by the LANCE® cAMP detection assay kit (PerkinElmer, Boston, MA, USA), in 384 well optiplates (PerkinElmer (Boston, MA, USA)) at 2000 cells/well in 5μl aliquots. Human αCGRP, hAM and hAM2 (Bachem, Switzerland) were diluted in PBS/BSA (0.1%) with 250μM IBMX (Sigma, UK), and used from 10pM to 10μM. Cells were incubated with compound for 30mins prior to adding detection buffer as described previously^28,57^. Plates were incubated for a further 60 mins (RT) and then read on a plate reader (Mithras LB 940 microplate reader (Berthold technologies, Germany)). All responses were normalised to 100μM forskolin (Tocris, UK). Antagonist studies were performed in the same way through co-stimulation of the relevant concentration. Alongside control treated cell. Experiments with PTX (Sigma, UK) required pretreatment (16hrs) prior to assays

#### Measurement of Phospho-ERK_1/2_ (Thr202/Tyr204)

Primary cells were grown to in 6 well plates, on the day of the experiment media was replaced with serum free media 4hrs prior to cell harvesting. Tyrpsin-EDTA was used to dissociate the cells and they are spun down, counted and re-suspended HBSS/BSA (0.1%). Ligands were also diluted in HBSS/BSA. Cells were then plated on 384 well plates in 8μl aliquots at a density of 20,000 cells/well. Next, ligands were added (4μl) for 5min stimulation at room temperature. Cells were then lysed as per to manufacturer’s instructions with 4μl of lysis buffer (Cisbio phosphor-ERK_1/2_ cellular assay kit, Invitrogen, UK) for 30mins shaking at room temperature. The 2 specific antibodies; were pre-mixed in a 1:1 ratio. 4μl of this was added to each well and the plate incubated for a further 2hrs. Then fluorescence emissions were read at 665nm and 620nm using a Mithras LB940 microplate reader. Antagonist studies were performed in the same way through co-stimulation with PTX, Rp-8-Br-cAMPS, YM-254890, or ESI-09 as appropriate alongside control treated cell.

#### Measurement of Intracellular Calcium mobilisation

All cell lines were plated at 20,000 cells/well on 96 well black clear-bottom plates (Costar, UK) 24hrs before the experiment. Media was removed, and cells were washed with Hank’s Balance Salt Solution (HBSS) (Lonza, Switzerland) before cells were loaded with 10μM Fluo-4/AM (Invitrogen, US) in the dark at room temperature for 30mins. Cells were then washed twice with calcium-free HBSS, then were left in 100μl calcium-free HBSS for the duration of the assay. In conditions where Gα_q/11/14_ signalling is inhibited, cells were pre-treated with 100nM YM-254890 (Alpha Laboratories, UK) (30mins)^22^. All assays were performed using the BD Pathway 855 Bioimaging Systems (BD Biosciences, UK), which dispenses ligands (20μl) and reads immediately for 2mins. Data was normalised to the response seen with 10μM Ionomycin.

#### Measurement of Cell Proliferation

Both endothelial cell lines and HCMs were seeded at a density of 2500 cells/well in a clear flat bottom 96-well plate (Corning, UK) and incubated at 37°C in 5% CO_2_. After 24 hrs, cells were exposed to test compounds or vehicle, in complete endothelial cell growth media (HUVECs) or myocyte growth media (HCMs). Cells were incubated for a further 72hrs at 37°C in 5% CO_2_. After 72 hrs incubation, 5μl of Cell Counting Kit – 8 (CCK-8, Sigma, UK) was added to each well and cells were then incubated for another 2 hrs at 37°C in 5% CO_2_ and in the dark. The absorbance of each well was measured using a Mithras LB940 microplate reader with an excitation of 450 nm. The absorbance is directly proportional to the number of viable cells. Cell proliferation was calculated as a percentage of number of cells treated with vehicle alone.

#### Measurement of Nitric Oxide Production

Endothelial cells and HCMs were cultured as above. 24 hours prior to assay cells were plated on Costar 96 well black clear bottom plates at 40,000 cells/well. The assay was performed according to manufacturer’s protocol. Briefly; cells pre-incubated with NO dye and assay buffer 1 (Fluorometric Nitric Oxide Assay Kit, Abcam, UK) for 30mins at 37°C in 5% CO_2_. Any inhibitors requiring 30mins pre-treatment (YM/L-NAME/DMSO control) were added at this point. Ligand stimulation occurred immediately after this for 15 mins at 37°C C in 5% CO_2_. Stain and ligand solution were removed, assay buffer II was added, and wells were read immediately. The absorbance was measured using a Mithras LB940 microplate reader with an excitation/emission of 540/590 nm. Endothelial cell responses were normalised to 10μM acetylcholine^58^. HCM responses were normalised to 10μM isoproterenol^59.60^.

## Statistical analysis

Data analysis for cAMP accumulation, Ca^2+^_i_ mobilisation, NO accumulation, pERK_1/2_ activation and cell proliferation assays were performed in GraphPad Prism 8.4 (GraphPad Software, San Diego). Data were fitted to obtain concentration–response curves using either the three-parameter logistic equation using to obtain values of E_max_ and pEC_50_ or the operational model of agonism^27^. Statistical differences were analysed using one-way ANOVA followed by Dunnett’s *post-hoc* (for comparisons amongst more than two groups) or unpaired Student’s t test with Welch’s correction (for comparison between two groups). To account for the day-to-day variation experienced from the cultured cells, we used the maximal level of cAMP accumulation from cells in response to 100μM forskolin stimulation was used as a reference, 10μM ionomycin for Ca^2+^_i_ assays, 10μM phorbol 12-myristate 13-acetate (PMA) for pERK_1/2_ activation, 10μM acetylcholine for NO production and 10μM VEGF for cell proliferation. E_max_ values from these curves are reported as a percentage of these controls, and all statistical analysis has been performed on these data. Where appropriate the operational model for receptor agonism^27^ was used to obtain efficacy (τ) and equilibrium disassociation constant (K_A_) values. In both cases, this normalization removes the variation due to differences in days but retains the variance for control values. The means of individual experiments were combined to generate the curves shown. Having obtain values for τ and K_A_ these were then used to quantify signalling bias as the change in Log(τ/K_A_) as described previously^18;24^. Error for this composite measure was propagated by applying the following equation.

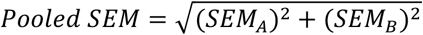

Where, *SEM*_*A*_ and *SEM*_*B*_ are the standard error of measurement A and B.

Correlations between pEC_50_ values or transducer coefficients Log (τ/K_A_) were assessed by scatter plot and Pearson’s correlation coefficient (*r*) was calculated with 95% confidence interval.

## Acknowledgements

This work was supported by the Biotechnology and Biological Sciences Research Council [grant number BB/M000176/2] awarded to GL and DRP and the Endowment Fund for education from Ministry of Finance Republic of Indonesia (DS). AJC is funded through an AstraZeneca Scholarship.

