## Supplementary material for "CGRP-receptor family reveals endogenous GPCR agonist bias and its significance in primary human cardiovascular cells": Supp Info

Supplementary Figures S1-S6
Supplementary Tables S1-S3

### Supplementary Figure 1

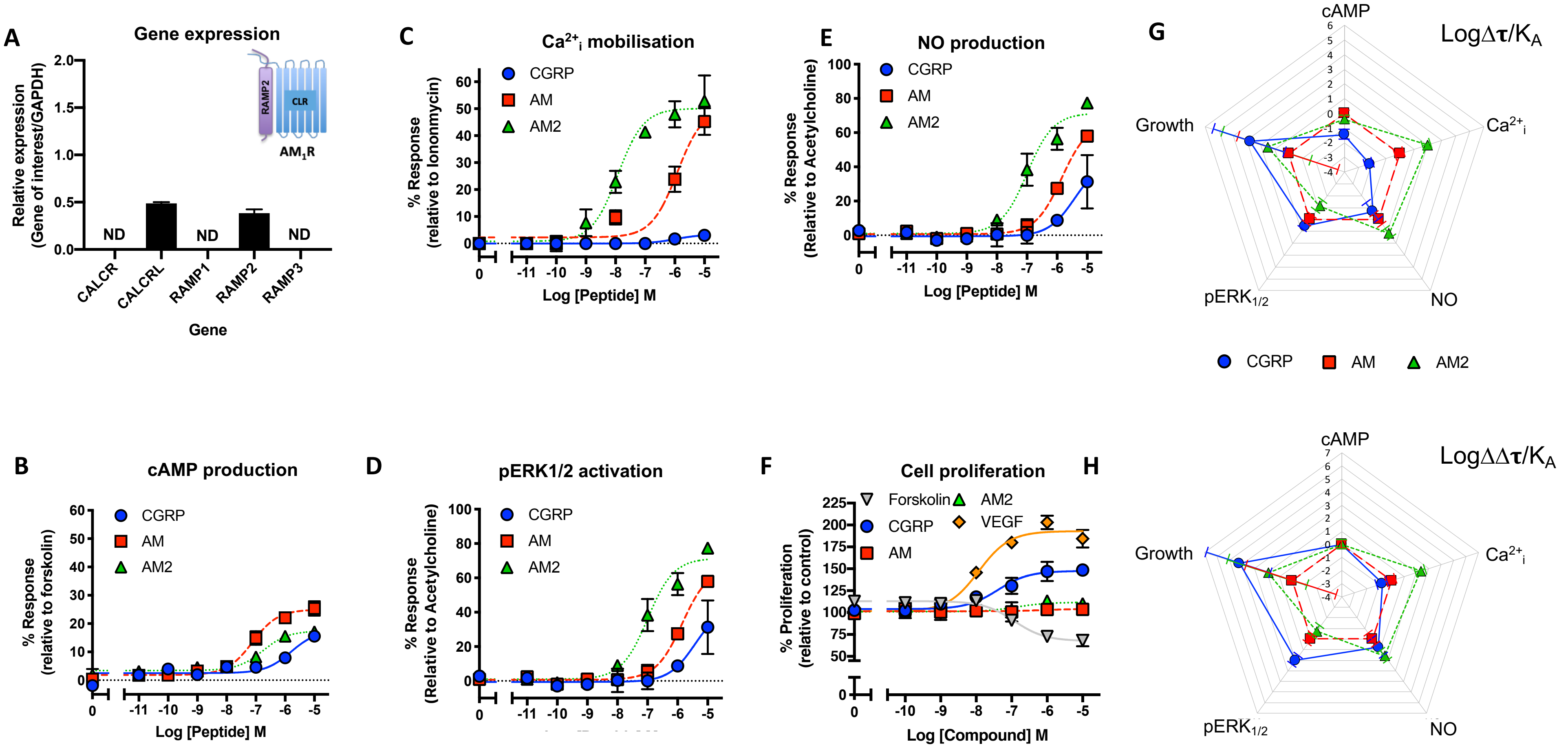


**Figure S1. CGRP family peptide signalling bias in HUAECs. A**) Expression of CALCR, CALCRL, RAMP1, RAMP2, and RAMP3 genes in HUVECs. Data represent mean + SEM of at least three independent experiments relative to GAPDH expression. ND = not detected in all three samples. **B-F)** Dose-response curves were constructed for HUVECs stimulated with CGRP, AM or AM2 and the cAMP levels quantified relative to forskolin (100μM) (**B**), mobilisation of $Ca^{2+}_i$ relative to ionomycin (10μM) (**C**), intracellular $ERK_{1/2}$ phosphorylation relative to PMA (10μM) (**D**), total NO production relative to acetylcholine (10μM) (**E**), and extent of cell proliferation (after 72 hours) relative to vector treated control and VEGF (**F**). Data are analysed using a three-parameter non-linear regression curve or the operational model of receptor agonism[27]. Data are analysed using a three-parameter non-linear regression curve. **G-H**) Signalling bias plots were calculated as $\Delta Log(\tau/K_A)$ (**G**) or $\Delta\Delta Log(\tau/K_A)$ (**H**) for each agonist and for each signalling pathway. Determination of values requires normalisation to a reference agonist (AM) alone in **G**, while for **H** values were normalised to both a reference agonist (AM) and a reference pathway (cAMP). All data represent mean ± SEM of at least 3-6 independent experiments.

**Supplemental Figure 2**

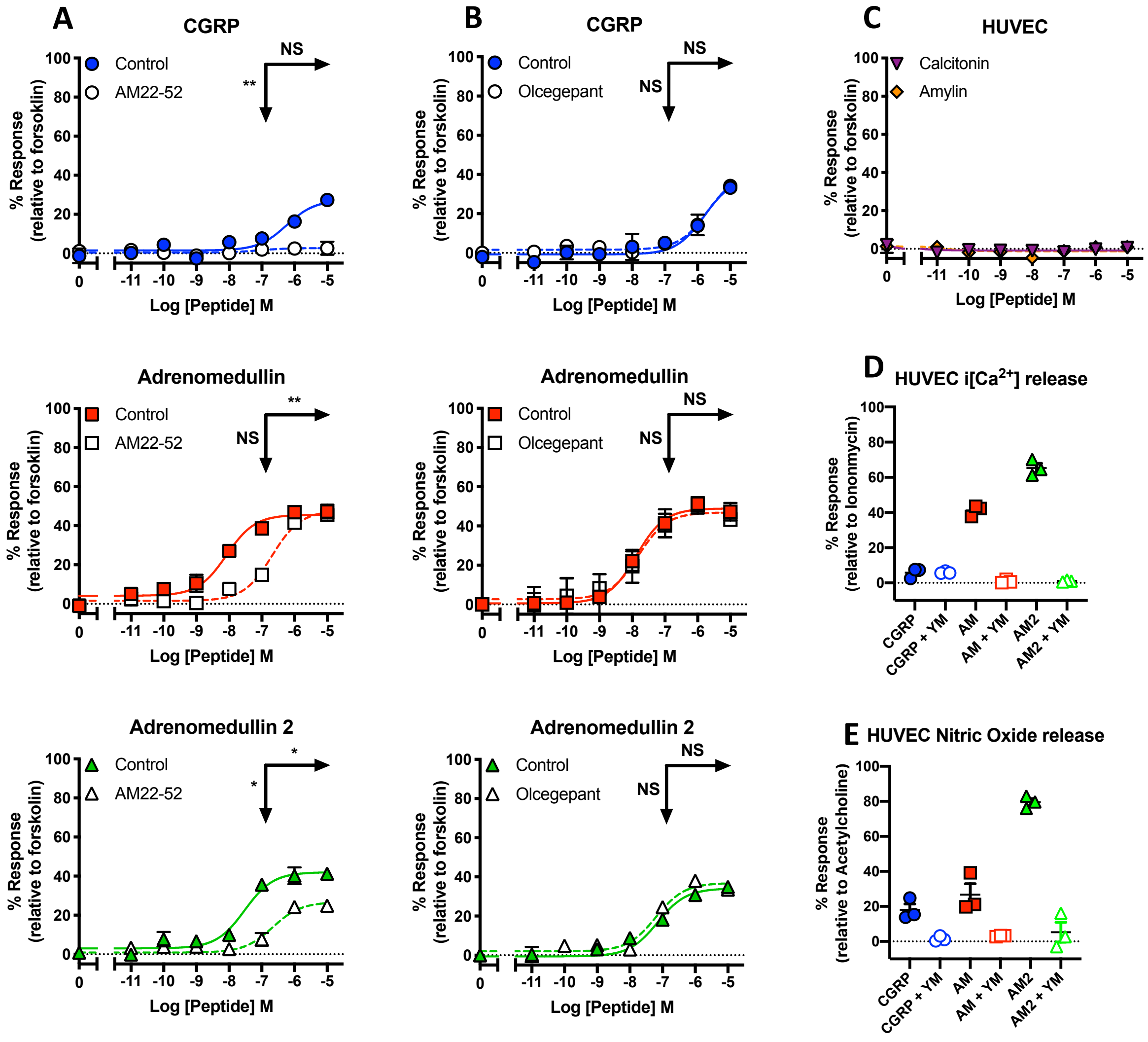


**Figure S2. Pharmacological assessment of signalling properties in HUVECs.**

**A-B**) Characterisation of cAMP accumulation in response to stimulation by CGRP, AM and AM2 in HUVECs in the presence and absence of 100nM AM22-52 (**A**) or 100nM Olcegepant (**B**). Data are analysed using a three-parameter non-linear regression curve. All data represent mean ± SEM of at least 3-6 independent experiments. Statistical significance determined compared to control using an unpaired Student's t test with Welch's correction (*, p<0.05; **, p<0.01; ***, p<0.001 ****, p<0.0001). NS denotes no statistical significance observed. Rows show $pEC_{50}$, and Vertical arrows show $E_{max}$ statistical significance. **C**) Characterisation of cAMP accumulation in response to stimulation by calcitonin and amylin in HUVECs relative to forskolin (100μM). **D**). Characterisation of $Ca^{2+}{}_i$ mobilisation with and without YM-254890 relative to ionomycin (10μM) and plotted as $E_{max}$ values. **E**) Characterisation of NO release with and without YM-254890 relative to Ach (10μM) and plotted as $E_{max}$ values ± SEM from at least 3 independent experiments.

### Supplemental Figure 3

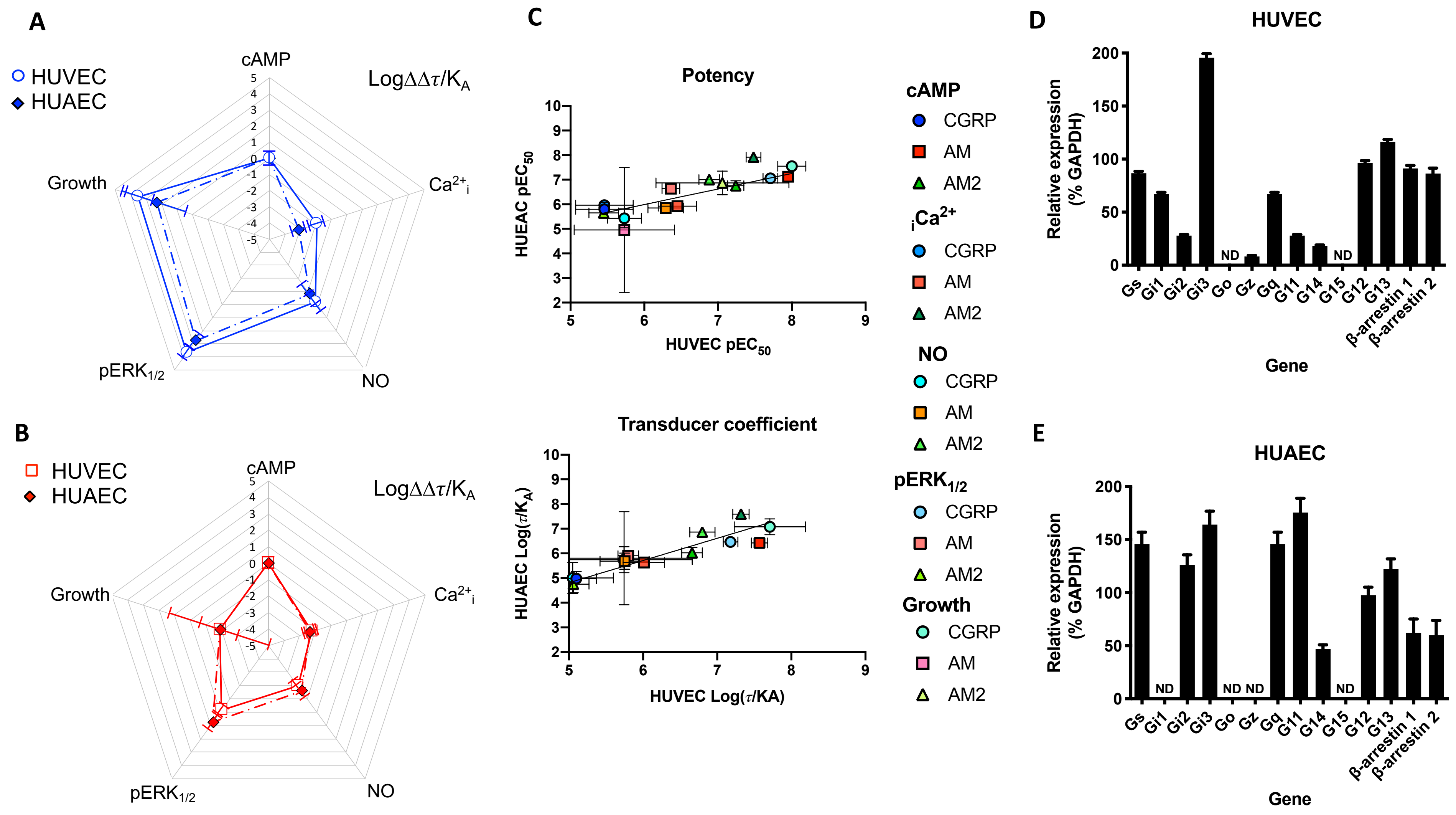


**Figure S3. Comparison of HUVEC and HUAEC signalling properties and G protein content**

**A**) Signalling bias plots were calculated as $\Delta\Delta\text{Log}(\tau/K_A)$ for CGRP in the two cell lines, HUVECs and HUAECs for each pathway. Values have been normalised to a reference agonist (AM2) and the reference pathway (cAMP) for both cell lines. **B**) as for **A** except the calculated values are for AM. **C**) the correlation of log agonist potencies ± SEM and transduction coefficient ($\tau/K_A$) ± SEM for CGRP, AM and AM2 stimulated cAMP accumulation, mobilisation of $Ca^{2+}_i$, NO production, intracellular $ERK_{1/2}$ phosphorylation and cell proliferation in HUVECs and HUAECs was analysed by a scatter plot and Pearson's correlation coefficients (*r*) were calculated. Significant positive correlation was observed for both potency and transduction coefficient between HUVECs and HUAECs as indicated by the presence of the line. **D-E)** Expression of GPCR accessory protein genes in HUVECs (**D**) and HUAECs (**E**). Data represent mean ± SEM of at least 3-6 independent experiments relative to GAPDH expression. ND = not detected in all three samples.

### Supplemental Figure 4

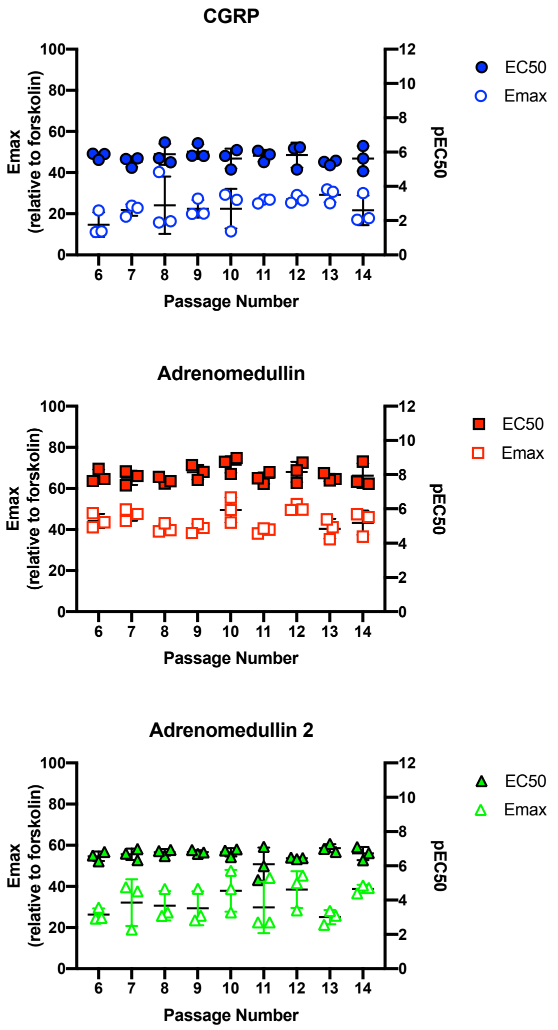


**Figure S4.**

**Quantification of CGRP signalling in HUVECs is unaffected by passage number** Characterisation of cAMP accumulation in response to stimulation by CGRP, AM and AM2 in HUVECs relative to 100μM forskolin across multiple passages (P6-P14). Data are analysed using a three-parameter non-linear regression curve, and $pEC_{50}/E_{max}$ values are plotted for at least 3 independent experiments ± SEM.

### Supplemental Figure 5

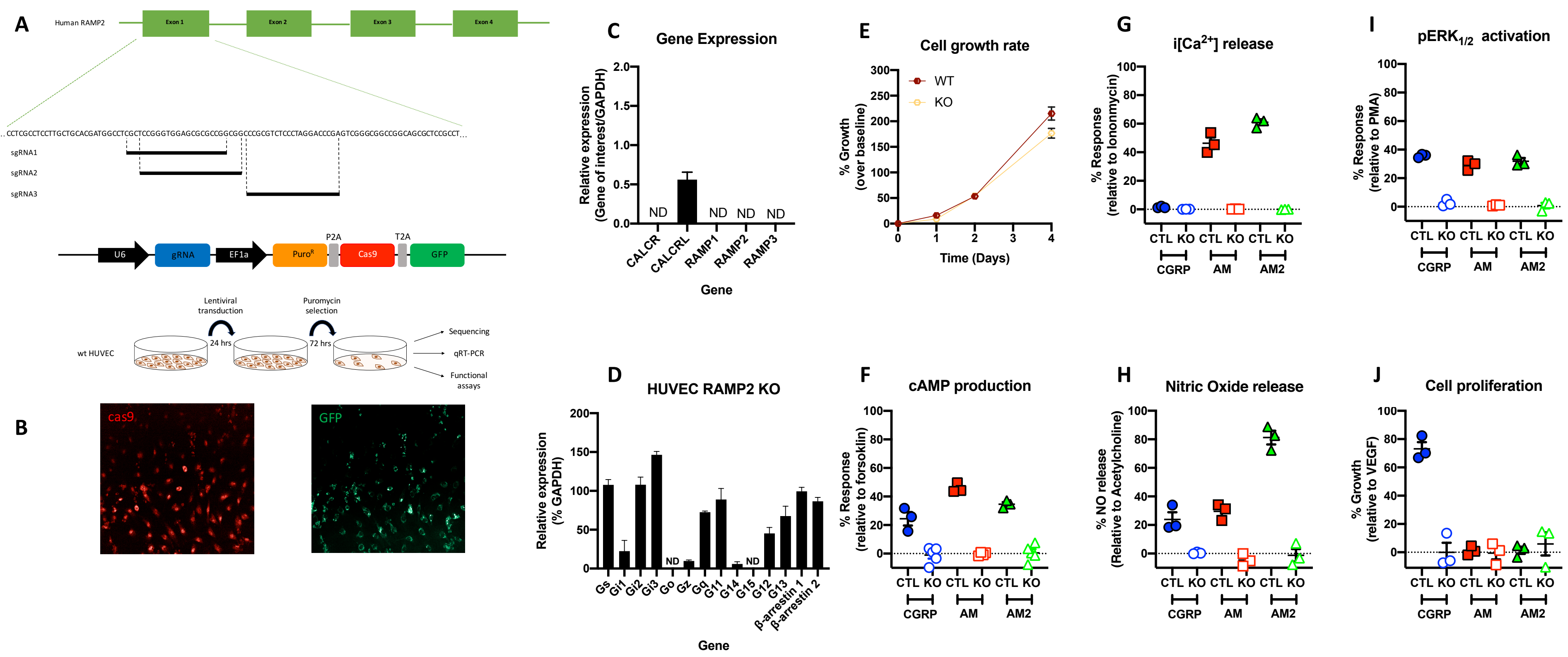


**Figure S5. Generation and confirmation of RAMP2 knockout HUVEC cell line by CRISPR-Cas9 gene editing and signalling characteristics and G protein content of HUVECΔRAMP2 cell line. A)** Schematic of the three gRNA sequences targeting the 1st exon of the WT RAMP2 gene and the lentiviral vector expressing gRNA, Puromycin resistance, GFP and Cas9 combined with the workflow for generation of gene edited HUVEC cell pool. **B)** Confocal imaging of HUVEC cells following transduction of lentivirus and puromycin selection. Showing high levels of Cas9 expression and nuclear localization and high levels of GFP expression with cytoplasmic expression. **C)** Expression of CALCR, CALCRL, RAMP1, RAMP2, and RAMP3 genes in. RAMP2 KO HUVECs. Data represent mean ± SEM of at least

3 independent experiments relative to GAPDH expression. ND = not detected in all three samples. **D**) Characterisation of cell growth rate in WT and RAMP2 KO HUVECs over 4 days relative to day zero. **E)** Expression of GPCR accessory protein genes in KO HUVECs. Data represent mean ± SEM of at least three independent experiments relative to GAPDH expression. ND = not detected in all three samples. **F-I**) WT and RAMP1 null HUVECs were stimulated with CGRP, AM and AM2 and the maximal response ($E_{max}$) quantified for **F**) cAMP accumulation relative to 100μM forskolin, **G**) $Ca^{2+}_i$ mobilisation relative to 10μM ionomycin, **H**) total NO production relative to 10μM acetylcholine, **I**) intracellular $ERK_{1/2}$ phosphorylation relative to 10μM PMA, **J**) cell proliferation relative to 10μM VEGF. All pathways measured in response to stimulation by CGRP, adrenomedullin (AM) and adrenomedullin 2 (AM2). All data represent $E_{max}$ ± SEM of 3-4 independent experiments.

### Supplemental Figure 6

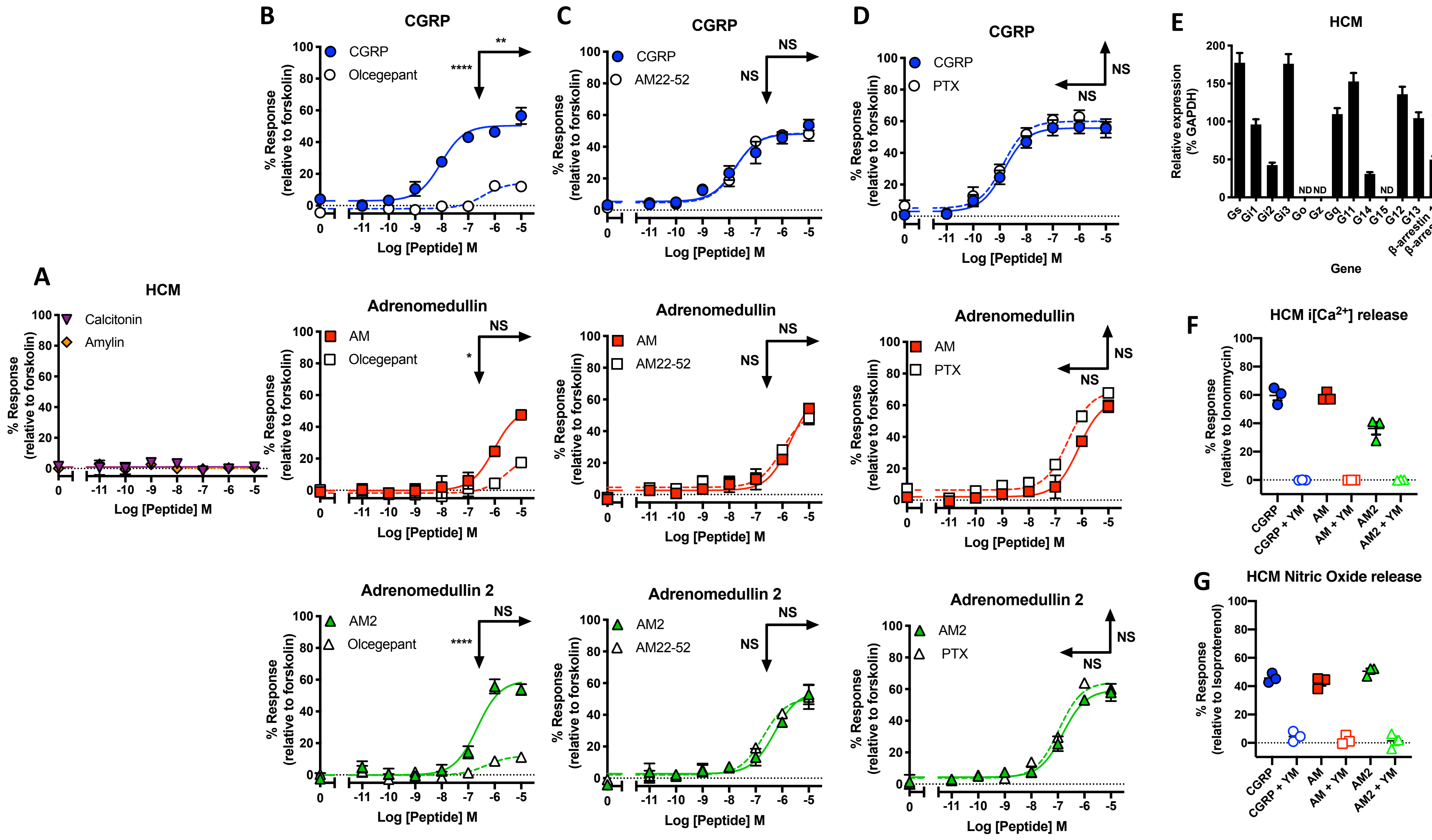

**Figure S6. Pharmacological assessment of signalling properties in HCMs.**

**A**) Characterisation of cAMP accumulation in response to stimulation by calcitonin and amylin in HCMs relative to forskolin (100μM). **B-C**) Characterisation of cAMP accumulation in response to stimulation by CGRP, AM and AM2 in HCMs in the presence and absence of 100nM Olcegepant (**B**) or AM22-52 (**C**). All relative to 100μM forskolin. **D**) Characterisation of cAMP accumulation in response to stimulation by CGRP, AM and AM2 with and without PTX treatment relative to forskolin (100μM) Data are analysed using a three-parameter non-linear regression curve. All data represent mean ± SEM of at least 3-6 independent experiments. Statistical significance determined compared to control using an unpaired Student's t test with Welch's correction (*, $p<0.05$; **, $p<0.01$; ***, $p<0.001$ ****, $p<0.0001$). NS denotes no statistical significance observed. All values are calculated from at least 3 individual data sets. Horizontal arrows show pEC50, and Vertical arrows show $E_{max}$ statistical significance. **E**) Expression of GPCR accessory protein genes in HCMs. Data represent mean ± SEM of at least three independent experiments relative to GAPDH expression. ND = not detected in all three samples. **F**) Characterisation of $Ca^{2+}{}_i$ mobilisation with and without YM-254890 relative to Ionomycin (10μM) and plotted as $E_{max}$ values. **G)** Characterisation of NO release with and without YM-254890 relative to acetylcholine (10μM) and plotted as $E_{max}$ values. Data in **F** and **G** represent $E_{max}$ ± SEM of 3-4 independent experiments.

**Table S1** Potency ($pEC_{50}$), effector Maximum ($E_{max}$), affinity ($pK_A$) and coupling efficacy (log τ) values for all signalling pathways various agonists measured in HUVECs and HUAECs.

| | HUVEC | | | HUAEC | | |
|---|---|---|---|---|---|---|
| | **cAMP** | | | | | |
| | **CGRP** | **AM** | **AM2** | **CGRP** | **AM** | **AM2** |
| **$pEC_{50}$[a]** | 5.46±0.39**** | 7.95±0.07 | 7.24±0.11 | 5.80±0.27*** | 7.12±0.16 | 6.75±0.20 |
| **$E_{max}$[b]** | 27.14±6.75 | 45.17±1.17 | 32.75±1.21 | 17.51±2.46* | 24.97±1.51 | 17.46±1.21* |
| **$pK_A$[c]** | 5.61±0.25**** | 7.67±0.09 | 6.99±0.14**** | 5.72±0.28*** | 6.95±0.14 | 6.78±0.21 |
| **logτ[d]** | -0.51±0.09*** | -0.10±0.07 | -0.33±0.03 | -0.74±0.09* | -0.52±0.03 | -0.75±0.05* |
| | **$Ca^{2+}_i$** | | | | | |
| **$pEC_{50}$[a]** | 5.45±0.33** | 6.47±0.26 | 7.48±0.10* | 5.96±0.14 | 5.92±0.17 | 7.92±0.14**** |
| **$E_{max}$[b]** | 2.26±0.54**** | 34.72±3.71 | 66.37±2.66**** | 3.36±0.29**** | 50.35±4.95 | 50.11±3.22 |
| **$pK_A$[c]** | 5.44±0.34 | 6.29±0.27 | 7.03±0.10 | 4.74±0.14 | 5.65±0.21 | 7.61±0.13**** |
| **logτ[d]** | -1.63±0.10**** | -0.28±0.07 | 0.29±0.05**** | -1.29±0.04**** | -0.01±0.08 | -0.01±0.05 |
| | **pERK1/2** | | | | | |
| **$pEC_{50}$[a]** | 7.71±0.10*** | 6.36±0.12 | 5.46±0.2** | 7.06±0.18 | 6.64±0.15 | 5.65±0.21* |
| **$E_{max}$[b]** | 35.17±1.13 | 30.16±1.75 | 33.30±4.59 | 24.43±1.62* | 16.48±0.94 | 17.59±2.44 |
| **$pK_A$[c]** | 7.46±0.10*** | 6.18±0.13 | 5.41±0.19** | 6.99±0.13 | 6.66±0.21 | 5.43±0.33* |
| **logτ[d]** | -0.28±0.02 | -0.38±0.04 | -0.35±0.08 | -0.52±0.03 | -0.75±0.05 | -0.67±0.12 |
| | **NO** | | | | | |
| **$pEC_{50}$[a]** | 5.73±0.23 | 6.29±0.24 | 6.88±0.14 | 5.43±0.39 | 5.85±0.09 | 7.00±0.18* |
| **$E_{max}$[b]** | 21.13±2.89 | 31.92±4.25 | 82.1±4.62*** | 43.25±13.02 | 66.07±2.63 | 71.48±4.88 |
| **$pK_A$[c]** | 5.64±0.52 | 6.08±0.31 | 6.15±0.15 | 5.10±0.56 | 5.41±0.30 | 6.47±0.14 |
| **logτ[d]** | -0.59±0.18 | -0.33±0.11 | 0.65±0.09** | -0.09±0.28 | 0.28±0.13 | 0.4±0.07 |
| | **Cell proliferation** | | | | | |
| **$pEC_{50}$[a]** | 8.00±0.19* | 5.73±0.68 | 7.06±0.9 | 7.55±0.19** | 4.96±2.54 | 6.87±0.48* |
| **$E_{max}$[b]** | 140.39±2.13**** | 106.90±3.47 | 103.74±1.43 | 147.55±3.07** | 119.22±4.44 | 111.68±1.62 |
| **$pK_A$[c]** | 7.69±0.19 | 5.69±0.70 | 7.03±0.90 | 7.43±0.31 | 5.30±1.80 | 6.91±1.87 |
| **logτ[d]** | 0.02±0.06** | -0.95±0.22 | -1.29±0.20 | -0.35±0.07 | -1.00±1.79 | -1.10±0.26 |

Data are the mean ± SEM of 3-6 individual data sets.
[a] The negative logarithm of the agonist concentration required to produce a half-maximal response.
[b] The maximal response to the agonist expressed as a percentage of the system maximal as define in the methods.
[c] The negative logarithm of the equilibrium disassociation constant for each ligand generated through use of the operational model of agonism[27].
[d] $\tau$ is the coupling efficiency parameter of each ligand.
Statistical significance compared to the cognate ligand AM (*, $p < 0.05$, **, $p < 0.01$, ***, $p < 0.001$, ****, $p < 0.0001$) for each pathway was determined by one-way ANOVA with Dunnett's post-test.

**Table S2** Potency ($pEC_{50}$), effector Maximum ($E_{max}$), affinity ($pK_A$) and coupling efficacy (log τ) values for all signalling pathways various agonists measured in RAMP1-HUVECs and HCMs.

| | RAMP1-HUVEC | | | HCM | | |
|---|---|---|---|---|---|---|
| | | | cAMP | | | |
| | **CGRP** | **AM** | **AM2** | **CGRP** | **AM** | **AM2** |
| **$pEC_{50}$[a]** | 9.03±0.12 | 7.43±0.17**** | 7.17±0.11**** | 8.39±0.12 | 5.84±0.08**** | 6.73±0.10**** |
| **$E_{max}$[b]** | 43.2±1.29 | 38.95±2.08 | 32.67±1.57** | 62.91±1.85 | 63.05±3.14 | 61.04±2.5 |
| **$pK_A$[c]** | 8.87±0.11 | 7.20±0.14**** | 6.97±0.16**** | 8.05±0.09 | 5.38±0.19**** | 6.29±0.14**** |
| **logτ[d]** | -0.15±0.02 | -0.22±0.03 | -0.34±0.05** | 0.20±0.03 | 0.22±0.10 | 0.18±0.06 |
| | | | **$Ca^{2+}_i$** | | | |
| **$pEC_{50}$[a]** | 7.90±0.12 | 7.24±0.09*** | 5.78±0.10**** | 9.30±0.10 | 8.25±0.17* | 8.17±0.33* |
| **$E_{max}$[b]** | 51.30±2.06 | 44.13±1.42* | 40.96±2.26** | 64.44±1.73 | 58.12±0.15 | 36.49±2.31*** |
| **$pK_A$[c]** | 7.52±0.09 | 6.97±0.10* | 5.64±0.18**** | 8.85±0.15 | 8.10±0.15 | 7.91±0.35 |
| **logτ[d]** | 0.03±0.03 | -0.10±0.03 | -0.18±0.06** | 0.22±0.05 | 0.08±0.04 | -0.30±0.05*** |
| | | | **pERK1/2** | | | |
| **$pEC_{50}$[a]** | 6.62±0.20 | 7.47±0.08* | 6.90±0.14 | 6.99±0.11 | 7.75±0.14* | 5.66±0.17** |
| **$E_{max}$[b]** | 38.69±3.4 | 47.57±1.48 | 45.35±2.89 | 33.02±1.52 | 40.13±2.02 | 40.41±4.17 |
| **$pK_A$[c]** | 6.52±0.17 | 7.16±0.13* | 6.56±0.14 | 6.77±0.15 | 7.52±0.12* | 5.52±0.18** |
| **logτ[d]** | -0.19±0.05 | -0.02±0.04 | -0.06±0.05 | -0.33±0.04 | -0.19±0.03 | -0.20±0.08 |
| | | | **NO** | | | |
| **$pEC_{50}$[a]** | 7.11±0.16 | 6.77±0.19 | 5.90±0.23** | 6.27±0.23 | 6.37±0.27 | 6.87±0.11 |
| **$E_{max}$[b]** | 75.15±4.51 | 56.30±4.5 | 53.96±7.60 | 49.97±5.56 | 46.98±5.85 | 50.59±2.18 |
| **$pK_A$[c]** | 6.54±0.18 | 6.43±0.21 | 5.51±0.29* | 5.99±0.22 | 6.07±0.22 | 6.55±0.18 |
| **logτ[d]** | 0.47±0.09 | 0.11±0.08 | 0.08±0.14 | 0.01±0.08 | -0.03±0.08 | 0.03±0.06 |
| | | | **Cell proliferation** | | | |
| **$pEC_{50}$[a]** | 6.07±0.49 | 6.77±0.40 | 5.85±0.17 | 5.32±0.80 | 7.08±0.23* | 6.48±0.58 |
| **$E_{max}$[b]** | 143.61±12.20 | 195.27±14.23* | 201.29±11.45* | 108.8±24.20 | 160.32±6.82* | 114.68±3.58 |
| **$pK_A$[c]** | 5.89±0.70 | 5.84±0.81 | 5.02±0.44 | 5.23±0.90 | 6.45±0.33 | 6.38±0.59 |
| **logτ[d]** | -0.09±0.30 | 1.10±0.64 | 0.71±0.34 | -0.65±0.80 | 0.52±0.20 | -0.60±0.12 |

Data are the mean ± SEM of 3-6 individual data sets.
[a] The negative logarithm of the agonist concentration required to produce a half-maximal response.
[b] The maximal response to the agonist expressed as a percentage of the system maximal as define in the methods.
[c] The negative logarithm of the equilibrium disassociation constant for each ligand generated through use of the operational model of agonism[27].
[d] $\tau$ is the coupling efficiency parameter of each ligand.
Statistical significance compared to the cognate ligand CGRP (*, $p < 0.05$, **, $p < 0.01$, ***, $p < 0.001$, ****, $p < 0.0001$) for each pathway was determined by one-way ANOVA with Dunnett's post-test.

**Table S3** Transducer coefficients Log($\tau$/$K_A$) as determined for all signalling pathways various agonists as measured in HUVECs, HUAECs, RAMP1-HUVEC and HCMs. Values calculated from the data shown in Table S1 and Table S2,

| | **CGRP** | **AM** | **AM2** |
|---|---|---|---|
| **HUVEC** | | | |
| **cAMP** | 5.10±0.27 | 7.57±0.11 | 6.66±0.14 |
| **$Ca^{2+}_i$** | 3.81±0.35 | 6.01±0.28 | 7.32±0.11 |
| **$pERK_{1/2}$** | 7.18±0.10 | 5.80±0.14 | 5.06±0.21 |
| **NO** | 5.05±0.55 | 5.75±0.33 | 6.80±0.17 |
| **Growth** | 7.71±0.48 | 4.74±0.73 | 5.74±0.92 |
| **HUAEC** | | | |
| **cAMP** | 4.98±0.29 | 6.43±0.14 | 6.03±0.22 |
| **$Ca^{2+}_i$** | 3.45±0.15 | 5.64±0.22 | 7.60±0.14 |
| **$pERK_{1/2}$** | 6.47±0.13 | 5.91±0.22 | 4.76±0.35 |
| **NO** | 5.01±0.63 | 5.69±0.33 | 6.87±0.16 |
| **Growth** | 7.08±0.32 | 4.30±2.54 | 5.81±1.89 |
| **RAMP1-HUVEC** | | | |
| **cAMP** | 8.72±0.11 | 6.98±0.14 | 6.63±0.17 |
| **$Ca^{2+}_i$** | 7.55±0.09 | 6.87±0.10 | 5.46±0.19 |
| **$pERK_{1/2}$** | 6.33±0.19 | 7.14±0.14 | 6.50±0.15 |
| **NO** | 7.01±0.20 | 6.54±0.22 | 5.59±0.32 |
| **Growth** | 5.80±0.76 | 6.94±1.03 | 5.73±0.56 |
| **HCM** | | | |
| **cAMP** | 8.25±0.09 | 5.60±0.21 | 6.47±0.15 |
| **$Ca^{2+}_i$** | 9.07±0.16 | 8.18±0.16 | 7.61±0.35 |
| **$pERK_{1/2}$** | 6.44±0.16 | 7.33±0.12 | 5.32±0.20 |
| **NO** | 6.00±0.23 | 6.04±0.23 | 6.58±0.19 |
| **Growth** | 4.58±1.20 | 6.97±0.39 | 5.78±0.60 |
